# The sterol-responsive RNF145 E3 ubiquitin ligase mediates the degradation of HMG-CoA reductase together with gp78 and Hrd1

**DOI:** 10.1101/391789

**Authors:** Sam A. Menzies, Norbert Volkmar, Dick J. van den Boomen, Richard T. Timms, Anna S. Dickson, James A. Nathan, Paul J. Lehner

## Abstract

HMG-CoA reductase (HMGCR), the rate-limiting enzyme of the cholesterol biosynthetic pathway and the therapeutic target of statins, is post-transcriptionally regulated by sterol-accelerated degradation. Under cholesterol-replete conditions, HMGCR is ubiquitinated and degraded, but the identity of the E3 ubiquitin ligase(s) responsible for mammalian HMGCR turnover remains controversial. Using systematic, unbiased CRISPR/Cas9 genome-wide screens with a sterol-sensitive endogenous HMGCR reporter, we comprehensively map the E3 ligase landscape required for sterol-accelerated HMGCR degradation. We find that RNF145 and gp78, independently co-ordinate HMGCR ubiquitination and degradation. RNF145, a sterol-responsive ER-resident E3 ligase, is unstable but accumulates following sterol depletion. Sterol addition triggers RNF145 recruitment to HMGCR and Insig-1, promoting HMGCR ubiquitination and proteasome-mediated degradation. In the absence of both RNF145 and gp78, Hrd1, a third UBE2G2-dependent ligase partially regulates HMGCR activity. Our findings reveal a critical role for the sterol-responsive RNF145 in HMGCR regulation and elucidate the complexity of sterol-accelerated HMGCR degradation.

## INTRODUCTION

Cholesterol plays a critical role in cellular homeostasis. As an abundant lipid in the eukaryotic plasma membrane, it modulates vital processes including membrane fluidity and permeability (Hannich et al., 2011; Haines, 2001) and serves as a precursor for important metabolites including steroid hormones and bile acids (Payne and Hales, 2004; Chiang, 2013). The cholesterol biosynthetic pathway in mammalian cells also provides intermediates for essential non-steroid isoprenoids and therefore requires strict regulation (Goldstein and Brown, 1990). The endoplasmic-reticulum (ER) resident, polytopic membrane glycoprotein 3-hydroxy-3-methylglutaryl coenzyme A reductase (HMGCR) is central to this pathway, catalysing the formation of mevalonate, a crucial isoprenoid precursor. As the rate-limiting enzyme in mevalonate metabolism, HMGCR levels need to be tightly regulated, as dictated by intermediates and products of the mevalonate pathway (Johnson and DeBose-Boyd, 2017). The statin family of drugs, which acts as competitive inhibitors of HMGCR, represents the single most successful approach to reducing plasma cholesterol levels and therefore preventing atherosclerosis related diseases (Heart Protection Study Collaborative Group, 2002). Understanding how HMGCR is regulated is therefore of fundamental biological and clinical importance.

Cholesterol, together with its biosynthetic intermediates and isoprenoid derivatives, regulates HMGCR expression at both the transcriptional and posttranscriptional level (Johnson and DeBose-Boyd, 2017). Low cholesterol induces transcriptional activation of HMGCR through the sterol response element binding proteins (SREBPs) which bind SREs in their promoter region (Osborne, 1991). In a cholesterol rich environment, SREBPs are inactive and held in the ER in complex with their cognate chaperone SREBP cleavage-activating protein (SCAP) in association with the ER-resident Insulin-induced genes 1/2 (Insig-1/2) anchor proteins (Dong and Tang, 2010; Yabe et al., 2002). A decrease in membrane cholesterol triggers dissociation of the SCAP-SREBP complex from Insigs and translocation to the Golgi apparatus, where the SREBP transcription factor is proteolytically activated by Site-1 and Site-2 proteases, released into the cytosol and trafficked to the nucleus (reviewed in Horton, Goldstein, & Brown, 2002). Low sterol levels therefore dramatically increase both HMGCR mRNA and extend HMGCR protein half-life, ensuring the resultant elevated enzyme levels stimulate the supply of mevalonate to re-balance cholesterol homeostasis (Goldstein and Brown, 1990; Brown et al., 1973). Once cholesterol levels are restored, excess HMGCR is rapidly degraded by the ubiquitin proteasome system (UPS) in a process termed sterol-accelerated degradation (Hampton et al., 1996; Ravid et al., 2000; Sever et al., 2003a). This joint transcriptional and translational regulation of HMGCR is therefore controlled by a host of ER-resident polytopic membrane proteins and represents a finely balanced homeostatic mechanism to rapidly regulate this critical enzyme in response to alterations in intracellular cholesterol. While the ubiquitin-mediated, post-translational regulation of HMGCR is well-established, the identity of the critical mammalian ER-associated degradation (ERAD) E3 ubiquitin ligase(s) responsible for sterol-accelerated HMGCR ERAD remains controversial.

In yeast, *S cerevisiae* encodes three ERAD E3 ligases, of which Hrd1p (HMG-CoA degradation 1), is named for its ability to degrade yeast HMGCR (Hmg2p) in response to non-sterol isoprenoids (Hampton et al., 1996; Bays et al., 2001). The marked expansion and diversification of E3 ligases in mammals makes the situation more complex, as in human cells there are 37 putative E3 ligases involved in ERAD, few of which are well-characterised (Kaneko et al., 2016). Hrd1 and gp78 represent the two mammalian orthologues of yeast Hrd1p. Hrd1 was not found to regulate HMGCR (Song et al., 2005; Nadav et al., 2003). However, gp78 was reported to be responsible for the sterol-induced degradation of HMGCR as (i) gp78 associates with Insig-1 in a sterol-independent manner, (ii) Insig-1 mediates a sterol-dependent interaction between HMGCR and gp78, (iii) overexpression of the transmembrane domains of gp78 exerted a dominant-negative effect and inhibited HMGCR degradation, and (iv), siRNA-mediated depletion of gp78 resulted in decreased sterol-induced ubiquitination and degradation of HMGCR (Song et al., 2005). The same laboratory subsequently suggested that the sterol-induced degradation of HMGCR was mediated by two ERAD E3 ligases, with TRC8 involved in addition to gp78 (Jo et al., 2011). However, these findings remain controversial as, despite confirming a for gp78 in the regulation of Insig-1 (Lee et al., 2006; Tsai et al., 2012), an independent study found no role for either gp78 or TRC8 in the sterol-induced degradation of HMGCR (Tsai et al., 2012). Therefore, the E3 ligase(s) responsible for the sterol-accelerated degradation of HMGCR remains disputed.

The introduction of systematic forward genetic screening approaches to mammalian systems (Carette et al., 2009; Wang et al., 2014) has made the unbiased identification of ubiquitin E3 ligases more tractable, as demonstrated for the viral (Greenwood et al., 2016; Van den Boomen and Lehner, 2015; van de Weijer et al., 2014; Stagg et al., 2009) and endogenous regulation of MHC-1 (Burr et al., 2011; Cano et al., 2012).

To identify the E3 ligases governing HMGCR ERAD we applied a genome-wide forward genetic screen to a dynamic, cholesterol-sensitive reporter cell line, engineered to express a fluorescent protein fused to endogenous HMGCR. This approach identified cellular genes required for sterol-induced HMGCR degradation, including UBE2G2 and the RNF145 ERAD E3 ligase. The subtle phenotype observed upon RNF145 depletion alone, suggested redundant ligase usage. A subsequent, targeted ubiquitome CRISPR/Cas9 screen in RNF145-knockout cells showed RNF145 to be functionally redundant with gp78, the E3 ligase originally implicated in HMGCR degradation. We confirmed that loss of gp78 alone showed no phenotype, while loss of both E3 ligases significantly inhibited the sterol-induced ubiquitination and degradation of HMGCR. Complete stabilisation required additional depletion of a third ligase - Hrd1. We find that endogenous RNF145 is an auto-regulated, sterol-responsive E3 ligase which is recruited to Insig proteins under sterol-replete conditions, thus promoting the regulated ubiquitination and sterol-accelerated degradation of HMGCR. Our data resolve the controversy of the E3 ligases responsible for the post-translational regulation of HMGCR and emphasise the complexity of the mammalian ubiquitin system in fine-tuning sterol-induced HMGCR turnover and cholesterol homeostasis.

## RESULTS

### Targeted knock-in at the endogenous HMMGCR locus creates a dynamic, cholesterol-sensitive reporter

To identify genes involved in the post-translational regulation of HMGCR, we engineered a cell line in which Clover, a bright fluorescent protein (Lam et al., 2012), was fused to the C-terminus of endogenous HMGCR, generating an HMGCR-Clover fusion protein (**Figure 1A**). The resulting HMGCR-Clover Hela single-cell clone expresses a dynamic, cholesterol-sensitive fluorescent reporter that is highly responsive to fluctuations in intracellular cholesterol. Basal HMGCR-Clover levels in sterol-replete tissue culture media were undetectable by flow cytometry (**Figure 1B**) and phenocopy endogenous WT HMGCR expression (**Figure 1C**, compare lanes 1 and 4). Following overnight sterol depletion, a ~ 25-fold increase in HMGCR-Clover expression was detected (shaded grey to blue histogram in **Figure 1D**, **Figure 1C** (lanes 2 and 5)), representing a combination of increased SREBP-induced transcription and decreased sterol-induced HMGCR degradation. Reintroduction of sterols induced the rapid degradation of HMGCR-Clover (~ 80% decrease within 2h), confirming the sterol-dependent regulation of the reporter (blue to red histogram in **Figure 1D**). Residual, untagged HMGCR detected by immunoblot in the reporter cells under sterol-depleted conditions suggested that at least one HMGCR allele remained untagged (**Figure 1C, compare lanes 2 and 5**). This unmodified allele allowed us to monitor both tagged and untagged forms of HMGCR. Inhibiting the enzymatic activity of HMGCR with mevastatin also stabilised HMGCR-Clover expression, as did inhibition of the proteasome (bortezomib) or p97 (NMS-873) (**Figure 1E**), confirming the rapid, steady-state degradation of the HMGCR reporter. Furthermore, we showed that CRISPR/Cas9-mediated ablation of both Insig-1 and -2 together induced a dramatic increase in HMGCR-Clover expression, equivalent to levels seen following sterol depletion (**Figure 1F**). Under these conditions, the SREBP-SCAP complex is not retained in the ER, leading to constitutive SREBP-mediated transcription of HMGCR-Clover, irrespective of the sterol environment. CRISPR-mediated gene disruption of either Insig-1 or -2 alone caused only a small, steady-state rescue of HMGCR-Clover (**Figure 1F**), which was more pronounced with the loss of Insig-1 than Insig-2. While Insig-1-deficient cells were unable to completely degrade HMGCR upon sterol addition, only a minor defect in HMGCR degradation was seen in the absence of Insig-2 (**Figure 1F**), suggesting that Insig-1 is dominant over Insig-2 under these conditions. Finally, we confirmed that HMGCR-Clover was appropriately localised to the ER by confocal microscopy (**Figure 1G**). Thus, HMGCR-Clover is a dynamic, cholesterol-sensitive reporter, which rapidly responds to changes in intracellular cholesterol and is regulated in a proteasome-dependent manner.

**Figure 1.**
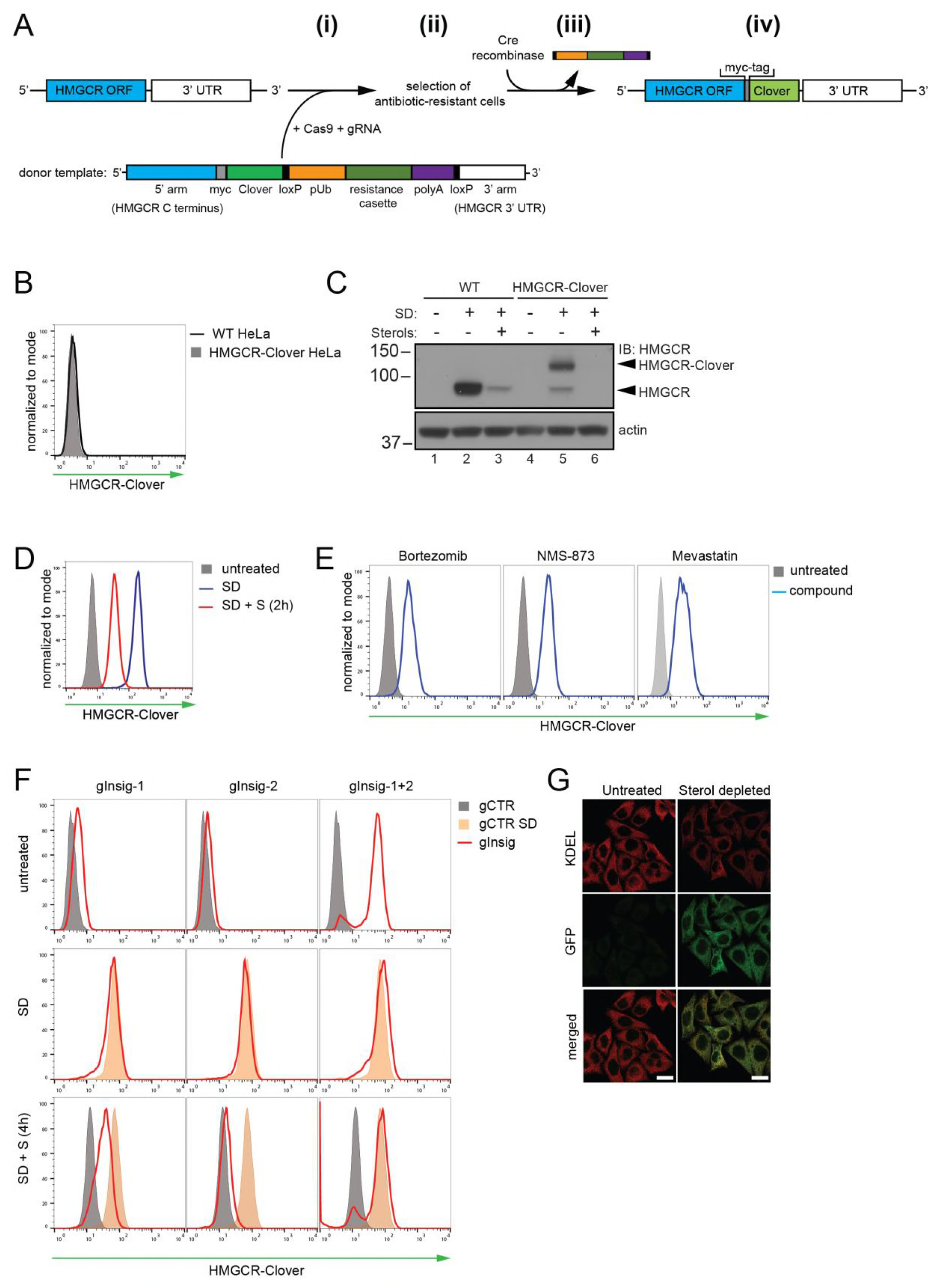
Fluorescent protein tagging of endogenous HMGCR generates a cholesterol-sensitive dynamic reporter. **(A)** Schematic showing generation of the HMGCR-Clover reporter. (i) The endogenous HMGCR locus of HeLa cells was modified by transfection of Cas9, gRNA and a donor template. The 5’ and 3’ arm of the donor template were designed as homologous sequences encoding the C-terminal region and 3’ UTR of the HMGCR gene. The C-terminal Clover (green) was appended in frame to the ORF of HMGCR (blue) including a myc-tag (grey) as spacer and an antibiotic resistance cassette flanked by loxP sites. (ii) Cells having stably integrated the recombination construct were enriched by antibiotic selection. (iii) The resistance cassette was removed by transient transfection of Cre recombinase to yield endogenous, C-terminally modified HMGCR (iv). ORF, open reading frame; UTR, untranslated region. **(B - E)** HMGCR-Clover reporter phenocopies untagged HMGCR. **(B)** HMGCR-Clover expression (shaded histogram) as detected by flow cytometry under sterol-replete conditions. **(C)** Immunoblot of HMGCR in sterol-depleted (SD) HeLa WT *vs*. HMGCR-Clover cells -/+ sterols (S) for 2h. For sterol depletion (SD), cells were switched to SD medium for 16h. Whole-cell lysates were separated by SDS-PAGE and HMGCR(-Clover) detected with an HMGCR-specific antibody. **(D)** Cytofluorometric analysis of HeLa HMGCR-Clover cells cultured in sterol-replete (shaded histogram) *vs*. sterol-depleted medium (SD) (16h, blue line histogram). Sterols (S) (2 μg/ml 25-hydroxycholesterol, 20 μg/ml cholesterol) were added back for 2h (red line histogram). **(E)** Flow cytometric analysis of HMGCR-Clover cells treated overnight with Bortezomib (25 nM), mevastatin (10 μM), or NMS-873 (10 μM) for 8h. **(F)** CRISPR/Cas9-mediated depletion of Insig-1 and -2 together induce a dramatic increase in HMGCR-Clover expression, equivalent to sterol depletion (SD). HMGCR-Clover cells transiently expressing the indicated Insig-1/2 specific gRNAs (4 sgRNAs per gene) were treated as in **(D)** and, where indicated, sterols (S) added back for 4h (SD + S, bottom row). Representative of ≥ 3 independent experiments. **(G)** Immunofluorescence analysis of HMGCR-Clover and KDEL (ER marker) expression, showing co-localisation in sterol-depleted (SD, 16h) cells. Scale bar = 20 μm.

### A genome-wide CRISPR/Cas9 screen identifies RNF145 as an E3 ligase required for HMGCR degradation

To identify genes required for the sterol-induced degradation of HMGCR, we performed a genome-wide CRISPR/Cas9 knockout screen in HMGCR-Clover cells. We took advantage of the rapid decrease in HMGCR-Clover expression following sterol addition to cells starved overnight (16h) of sterols (**Figure 1D**), and enriched for rare genetic mutants with reduced ability to degrade HMGCR-Clover in response to sterols. To this end, HMGCR-Clover cells were transduced with a genome-wide CRISPR/Cas9 knockout library comprising 10 sgRNAs per gene (Morgens et al., 2017). Mutagenised cells were first depleted of sterols overnight; sterols were then reintroduced for 5h, at which point rare mutant cells with reduced ability to degrade HMGCR-Clover upon sterol repletion were enriched by fluorescence-activated cell sorting (FACS) (**Figure 2A**, gating shown in **Figure 2 – figure supplement 1A**). This process was repeated again eight days later to further purify the selected cells. The enriched population contained only a small percentage of cells (1.96% after sort 1, 24.49% after sort 2) with increased steady-state HMGCR-Clover expression (green filled histogram in **Figure 2B**). However, the majority of sterol-starved cells from this selected population showed impaired degradation of HMGCR-Clover after addition of sterols (compare red *versus* orange histogram (**Figure 2B**, compare lanes 6 and 9 in **Figure 2 – figure supplement 1B)**). The broad distribution of this histogram (**Figure 2B** red histogram) suggested that the enriched cell population contains a variety of mutants which differ in their ability to degrade HMGCR-Clover.

**Figure 2.**
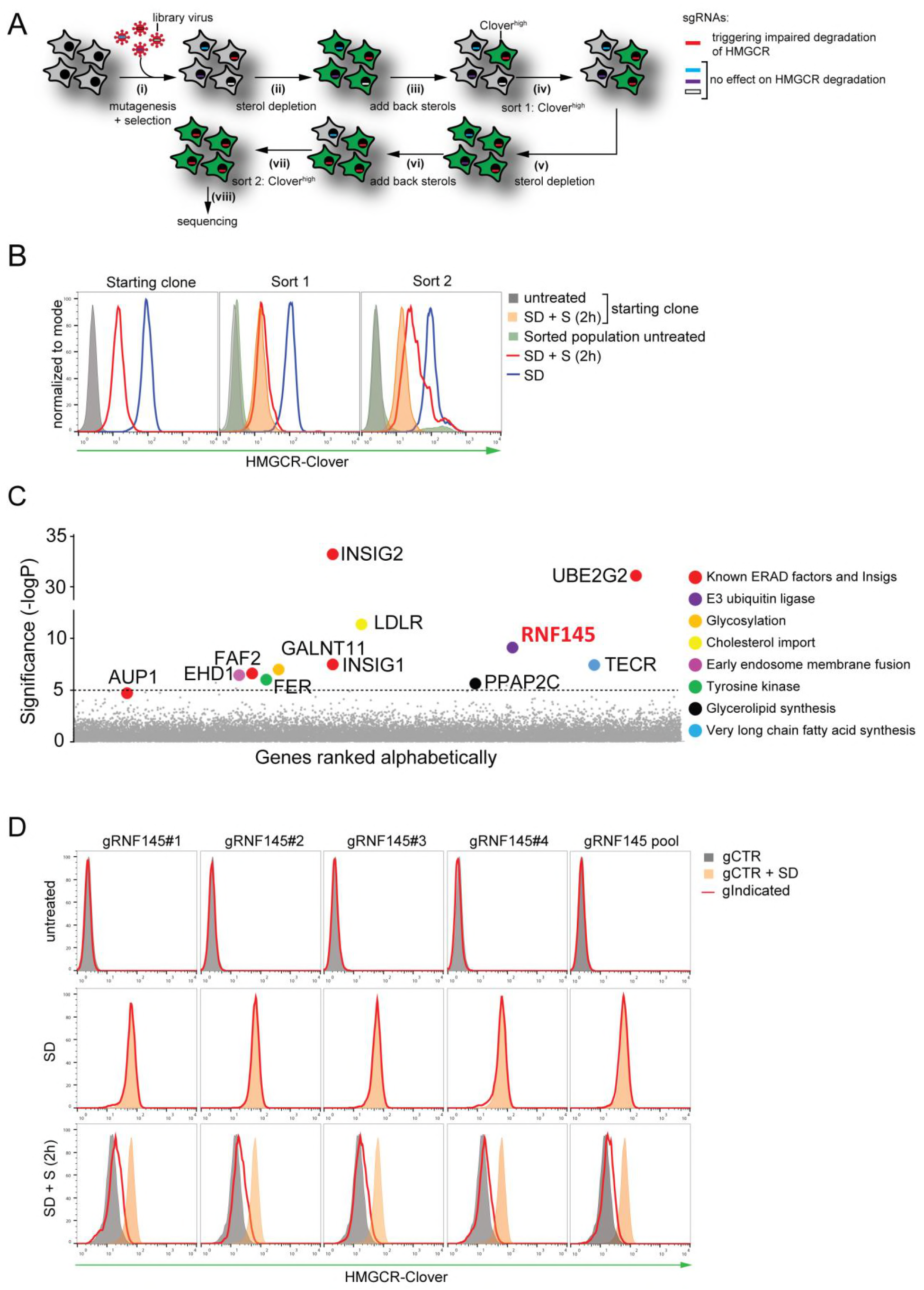
Genome-wide CRISPR knockout screen identifies a role for RNF145 in the sterol-dependent degradation of HMGCR. **(A - B)** Schematic view of the CRISPR/Cas9 knockout screen workflow and FACS enrichment. **(A)** HMGCR-Clover cells transduced with a genome-wide gRNA library targeting 19930 genes (i) were subjected to sterol-starvation/repletion (ii, iii, v, vi). Mutants unable to degrade HMGCR-Clover despite sterol repletion (Clover^high^) were enriched by two sequential rounds of FACS (iv, vii) and candidate genes identified by deep sequencing (viii). **(B)** Enrichment of HMGCR-Clover mutants after sort 1 and sort 2 (red line histograms, corresponding to steps ‘iv’ and ‘vii’ in **Figure 2A**) as determined by flow cytometry. Cells were treated as described in **Figure 1D**. SD, sterol-depleted; S, sterols. **(C)** Candidate genes identified in the genome-wide knockout screen. Genes scoring above the significance threshold of −logP ≥ 5 (dotted line) and AUP1 (- logP = 4.7) are highlighted. **(D)** RNF145 depletion mildly impairs sterol-accelerated HMGCR-Clover degradation. HMGCR-Clover cells transiently expressing 4 independent RNF145-specific sgRNAs (gRNF145#1-4, red line histogram), individually or as a pool, *vs*. gB2M (gCTR) were sterol-depleted overnight (middle row, SD) and re-examined by flow cytometry following 2h sterol addition (bottom row, SD + S). Representative of ≥ 3 independent experiments.

The sgRNAs in the selected cells, and an unselected control library, were sequenced on the Illumina HiSeq platform (**Figure 2A (viii)**). Using the RSA algorithm, we identified a set of 11 genes (-logP > 5), which showed significant enrichment in the selected cells. Many of these are known to be required for the sterol-induced degradation of HMGCR (**Figure 2C**) (König et al. 2007). The screen identified the E2 conjugating enzyme *UBE2G2* and its accessory factor *AUP1*, which recruits UBE2G2 to lipid droplets and membrane E3 ubiquitin ligases (Klemm et al., 2011; Jo et al., 2013; Spandl et al., 2011; Christianson et al., 2012), as well as both Insig-1 and -2 (Yabe et al., 2002; Yang et al., 2002; Sever et al., 2003a). The role of the remaining hits is summarized (**Figure 2 – figure supplement 2**) and validation of selected hits as shown (Insig-1/2, **Figure 1D**; UBE2G2, EHD1, GALNT11, LDLR and TECR, **Figure 2 – figure supplement 1C/D**).

Strikingly, the only ER-resident ubiquitin E3 ligase to emerge from the screen is RNF145, a poorly characterised ER-resident ubiquitin E3 ligase. RNF145 shares 27% amino acid identity with TRC8, which is one of the E3 ligases (together with gp78) previously suggested to ubiquitinate HMGCR (Jo et al., 2011). Interestingly, RNF145 also harbours a YLYF motif at its N-terminus, which is similar to the YIYF motif present in the sterol-sensing domain (SSD) of SCAP and HMGCR required for their binding to the Insig proteins (Yang et al., 2002; Sever et al., 2003a; Jiang et al., 2018; Cook et al., 2017; Zhang et al., 2017). The presence of the YLYF motif suggested that RNF145 might itself interact with the Insig proteins and therefore represented a promising candidate from our genetic screen.

To examine the role of RNF145 in HMGCR degradation, we designed four independent sgRNAs, either targeting RNF145 individually or as a pool. Under cholesterol-replete conditions, no accumulation of the HMGCR-reporter was observed in RNF145-depleted cells (top and middle rows, **Figure 2D**), but a small and highly reproducible decrease in HMGCR-Clover degradation was seen following re-introduction of sterols (red histograms, bottom row **Figure 2D**), emphasising the utility of the endogenous fluorescent reporter in identifying subtle phenotypes. Since the identity of the E3 ligases regulating HMGCR turnover remains controversial, the modest effect of RNF145 loss on HMGCR-Clover sterol-induced degradation suggested the involvement of additional ligase(s). Our screen therefore identified both known and novel components implicated in sterol-dependent HMGCR ERAD.

### RNF145 together with gp78 are required for HMGCR degradation

If a second E3 ligase is partially redundant with RNF145, its effect should be unmasked in RNF145-deficient cells. We therefore generated a focussed subgenomic sgRNA library targeting 1119 genes of the ubiquitin-proteasome system as described in **Materials and Methods**, including 830 predicted ubiquitin E3 ligases, and used this library to screen for genes required for the degradation of HMGCR in RNF145-deficient HMGCR-Clover cells (**Figure 3 – figure supplement 4B**, lane 2 for knockout validation). Due to the reduced complexity of this focussed library, only a single FACS enrichment step was used (**Figure 3A**, red histogram).

**Figure 3.**
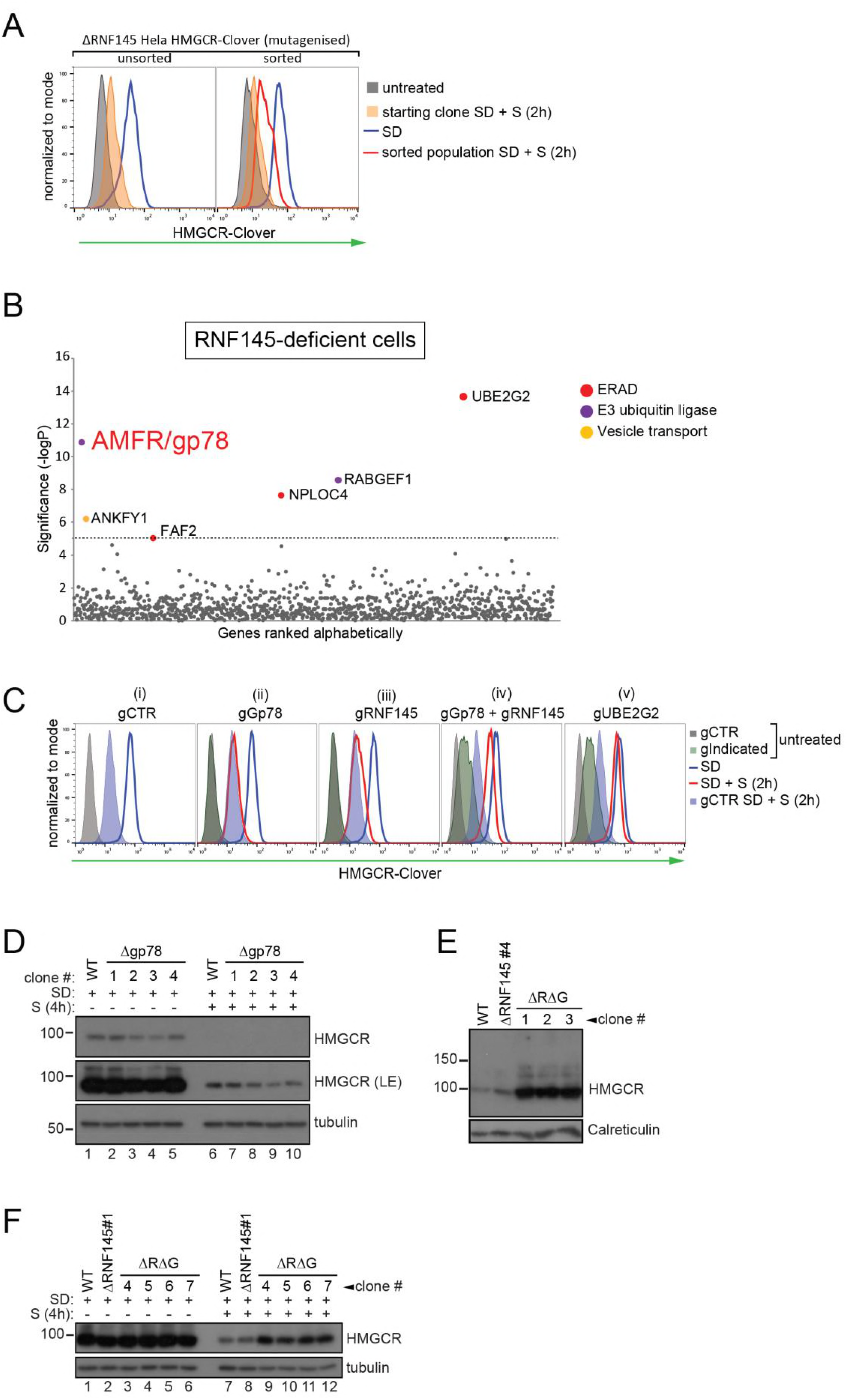
RNF145 together with gp78 are required for HMGCR degradation. **(A - B)** FACS enrichment and scatter plot of candidate genes identified in the ubiquitome-targeted knockout screen. **(A)** HeLa HMGCR-Clover ΔRNF145#5 cells were mutagenized using a targeted ubiquitome-specific sgRNA library and mutant cells showing impaired sterol-dependent degradation of HMGCR-Clover were enriched by FACS. Enrichment is represented by a broad population of Clover^high^ cells in the presence of sterols (S, 2h) after overnight sterol depletion (blue to red histogram). **(B)** Genes scoring above the significance threshold of - logP ≥ 5 (dotted line) are highlighted. **(C - F)** sgRNA targeting of gp78 together with RNF145 increases steady-state HMGCR-Clover and inhibits sterol-accelerated degradation of sterol-starved HMGCR-Clover. **(C)** HMGCR-Clover cells transiently transfected with indicated sgRNAs were sterol-depleted (SD) overnight (blue line histogram) and sterols (2 μg/ml 25-hydroxycholesterol, 20 μg/ml cholesterol) added back (SD+S) for 2h (red line histogram or blue shaded histogram for gCTR). Representative of ≥ 3 independent experiments. **(D)** Four independent gp78 knockout clones (#1-4) or WT cells were sterol-depleted (16h) ± S (4h) and HMGCR levels monitored by immunoblotting. LE, long exposure. **(E)** HMGCR steady-state levels in three RNF145/gp78 double knockout clones (ΔRΔG #13). **(F)** Four RNF145/gp78 double-knockout clones (ΔRΔG #4 - 7), RNF145 knockout, and WT cells were sterol-depleted (SD) overnight and HMGCR expression assessed ± sterols (4h) by immunoblot analysis.

Strikingly, this screen identified gp78 (gene name: *AMFR*) (**Figure 3B**, **Figure 3 – figure supplement 5**), the E3 ligase previously implicated in HMGCR degradation (Jo et al., 2011; Song et al., 2005; Fang et al., 2001). Taking a combined knockout strategy we asked whether gp78 and RNF145 are together responsible for HMGCR degradation. As predicted by the genetic approach (**Figure 3C(ii)**), there was no difference in sterol-induced HMGCR-Clover degradation between control and gp78-depleted HMGCR-Clover cells. Gp78 was not, therefore, a false-negative from our initial, genome-wide CRISPR/Cas9 screen (**Figure 2C**). Individual knockout of RNF145 again showed that sterol-induced HMGCR-Clover degradation was mildly impaired in RNF145-depleted cells (**Figure 3C (iii)**). However, sgRNA-mediated targeting of gp78 together with RNF145 (**Figure 3C (iv)**, see **Figure 3 – figure supplement 4A** and **B** lane 3 for knockout validation), resulted in a significant increase in both steady-state HMGCR-Clover (**Figure 3C (iv)** grey to green filled histograms) and an inability to degrade HMGCR-Clover upon addition of sterols to sterol-starved cells (**Figure 3C (iv)** blue to red histogram), a phenotype comparable to UBE2G2 deletion (**Figure 3C (v)**). Our results therefore suggest a partial functional redundancy between gp78 and RNF145 and imply that both ligases can independently regulate the sterol-induced degradation of HMGCR.

### RNF145 and gp78 regulate endogenous wild type HMGCR

To confirm that the phenotypes observed in RNF145- and gp78-deficient HMGCR-Clover cells were representative of endogenous, wild type HMGCR regulation, we deleted RNF145 and/or gp78 from WT HeLa cells and monitored endogenous HMGCR by immunoblot analysis. The sterol-induced degradation of HMGCR was assessed in four RNF145 knockout clones, derived from two different sgRNAs (validation in **Figure 3 – figure supplement 1A, B**). No difference in the sterol-induced degradation of HMGCR was seen in these RNF145 knockout clones (**Figure 3 – figure supplement 2**, compare lanes 6 and 7-10). The subtle effect on HMGCR-Clover expression revealed by flow cytometry (**Figures 2D and 3C**) may not be detected by the less sensitive immunoblot analysis. Similarly, loss of gp78 alone (**Figure 3 – figure supplement 3A** for sgRNA validation) did not affect HMGCR degradation (**Figure 3D**, compare lanes 6 and 7-10), but loss of gp78 together with RNF145 resulted in a significant rescue of steady state HMGCR (**Figure 3E**, **Figure 3 – figure supplement 3C**). Following sterol addition, gp78/RNF145 double-knockout clones showed a marked (although still incomplete) reduction in sterol-induced HMGCR degradation (**Figure 3F**, compare lanes 7+8 with 9-12). These data validate the phenotypes exhibited by the HMGCR-Clover reporter cell line and confirm a role for both gp78 and RNF145 in the sterol-induced degradation of endogenous HMGCR.

### RNF145 E3 ligase activity is required for HMGCR degradation

To determine whether RNF145 E3 ligase activity is required for HMGCR degradation, we complemented a mixed population of gp78/RNF145 double-knockout HMGCR-Clover cells (**Figure 3 – figure supplement 4B**, lane 3 for knockout validation) with either epitope-tagged wild type RNF145, or a catalytically-inactive RNF145 RING domain mutant (C552A, H554A) (**Figure 4A**). The pronounced block in the sterol-induced degradation of HMGCR-Clover was rescued by expression of wild-type, but not the RNF145 RING domain mutant (**Figure 4B**, compare blue to red histogram). The E3 ligase activity of RNF145 is therefore critical for HMGCR ERAD.

**Figure 4.**
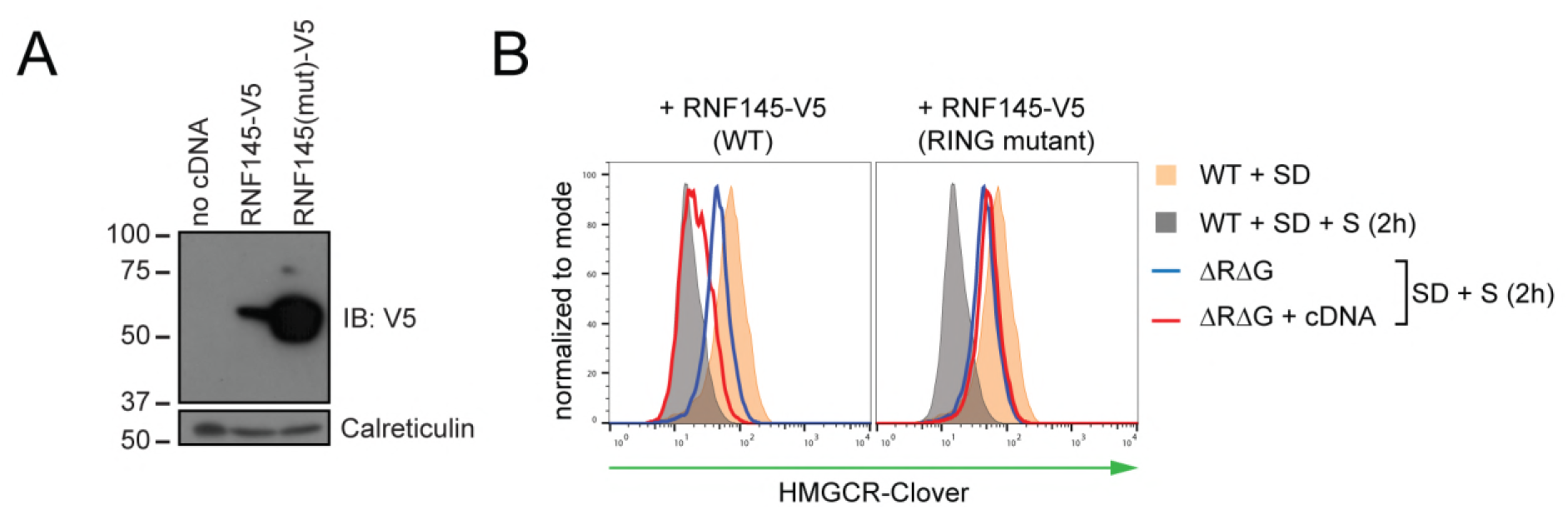
RNF145 E3 ligase activity is required for HMGCR degradation. **(A)** Exogenous expression of RNF145 and RING-mutant RNF145 in HMGCR-Clover cells. RNF145/gp78 double-knockout HMGCR-Clover cells were transduced with lentivirus expressing either RNF145-V5 or a catalytically inactive RING domain mutant RNF145(C552A, H554A)-V5 cDNA and cell lysates separated by SDS-PAGE and subsequent immunoblot analysis. IB, immunoblot. **(B)** Wildtype (WT) but not RING mutant RNF145 complements the RNF145-deficient phenotype. RNF145/gp78 double-knockout HMGCR-Clover cells (ΔRΔG #11) were transduced with lentivirus expressing either RNF145-V5 or a catalytically inactive RING domain mutant RNF145(C552A, H554A)-V5 cDNA. Cells were sterol-depleted (16h) and after sterol repletion (2h), HMGCR-Clover levels were assessed by flow cytometry.

### Endogenous RNF145 is an unstable E3 ligase, whose transcription is sterol-regulated

Endogenous RNF145 has a short half-life (~ 2h) and displayed rapid, proteasome-mediated degradation (**Figure 5A (i)**), an observation confirmed in multiple cell lines (**Figure 5 – figure supplement 1A**). This rapid turnover of endogenous RNF145 contrasts sharply with the stability of endogenous gp78, which shows little degradation over the 10 hour chase period (**Figure 5A (i)**). Although RNF145 and gp78 both target HMGCR for degradation, the two ligases did not appear to be co-regulated as RNF145 stability was unaffected by gp78 and vice-versa (**Figure 5A (i, ii)**, **Figure 5 – figure supplement 1B**). However, endogenous RNF145 was stabilised by deletion of its cognate E2 enzyme UBE2G2 (**Figure 5B**), and, furthermore, the catalytically-inactive RING domain mutant expressed in RNF145-deficient cells (ΔR145#4 + R145-V5 (mut)) exhibited greater abundance at steady-state compared with its wild type counterpart (**Figure 3 – figure supplement 4C**). Together these data show that RNF145 is intrinsically unstable and rapidly turned over in an auto-regulatory manner.

**Figure 5.**
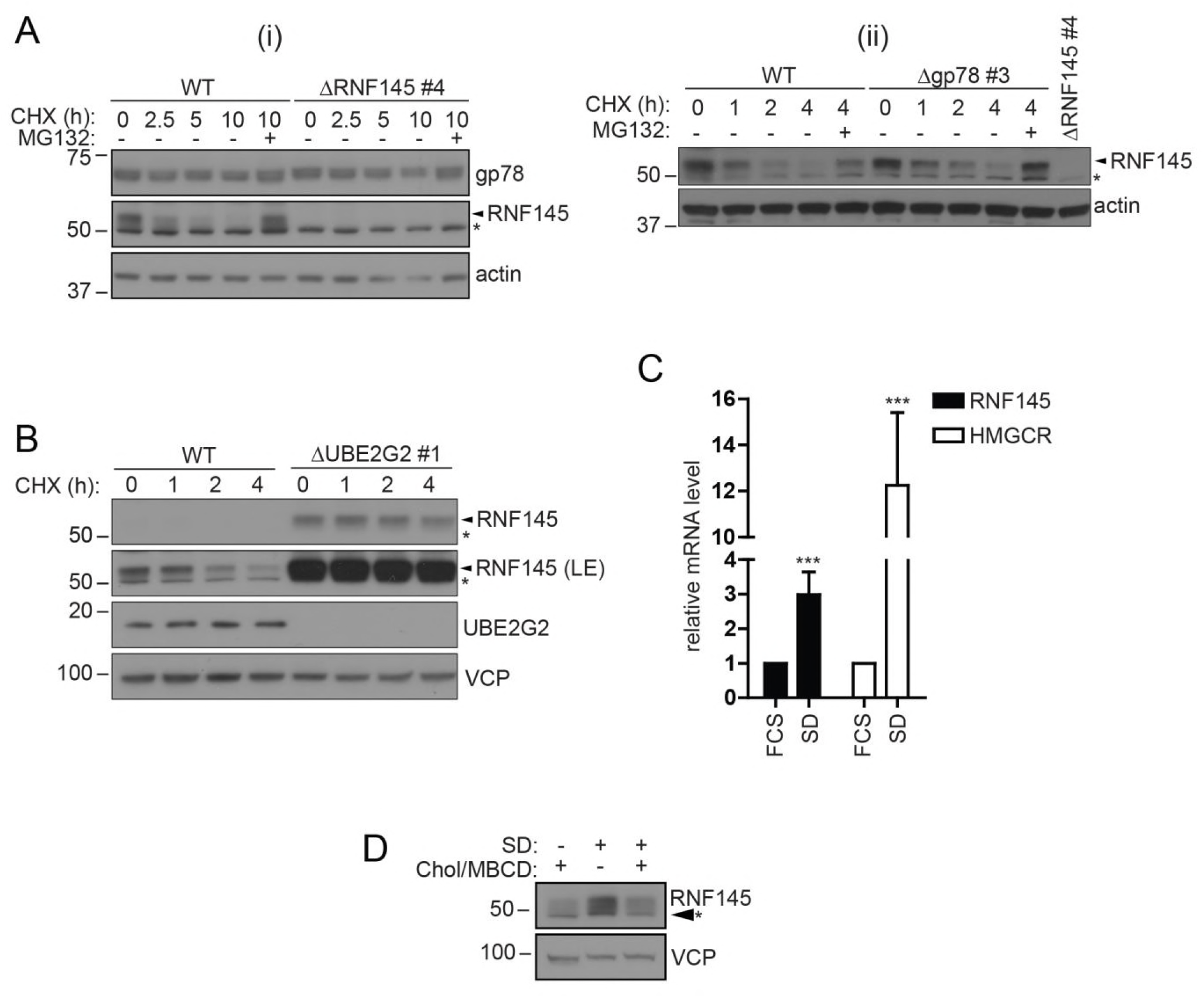
RNF145 is an intrinsically unstable, sterol-responsive E3 ligase. **(A and B)** RNF145 has a short half-life and is auto-regulated by UBE2G2. **(A)** Translational shutoff analysis of gp78 in WT *versus* ΔRNF145 #4 (i) or RNF145 in Δgp78 #3 cells (ii) treated with cycloheximide (CHX, 1 μg/ml) ± MG-132 (20 μg/ml) for the indicated times. Non-specific bands are indicated by an asterisk (*). Representative of ≥ 2 independent experiments. **(B)** Immunoblot analysis of WT and ΔUBE2G2 cells treated with CHX (1 μg/ml) for the indicated times. VCP serves as a loading control. LE, long exposure. **(C and D)** Sterol depletion induces transcriptional activation and increased levels of RNF145 protein. **(C)** Relative RNF145 and HMGCR mRNA levels as measured by qRT-PCR in HeLa cells grown in 10% FCS (FCS) or sterol-depleted (SD, 10% LPDS + 10 μM mevastatin + 50 μM mevalonate) for 48 h. Mean ± S.D. (n = 4) and significance are shown, unpaired Students t-test: ***p ≤ 0.001. **(D)** HeLa cells were grown under sterol-rich or sterol-deplete conditions (as indicated) for 48 h in the presence of mevastatin (10 μM) and mevalonate (50 μM) ± complexed cholesterol (chol:MBCD, 37.5 μM). Whole cell lysates were separated by SDS-PAGE and underwent immunoblot analysis. Non-specific bands are indicated (*). Representative of ≥ 2 independent experiments.

Since RNF145 is rapidly turned over, we aimed to determine whether RNF145 gene transcription was sterol-responsive. Sterol depletion induced RNF145 (~2.99±0.65 fold increase, p = 0.0009) mRNA expression as well as HMGCR (~12.26±3.16 fold increase, p = 0.0004) mRNA expression (**Figure 5C**). This accumulation of endogenous RNF145 was suppressed following the addition of MBCD-complexed cholesterol (chol:MBCD) to the starvation media (**Figure 5D**), whereas gp78 abundance remained unaltered (**Figure 5 – figure supplement 1D**). RNF145 is therefore a unique, sterol-regulated E3 ligase whose expression is dependent on the cellular sterol status.

### Endogenous RNF145 shows a sterol-sensitive interaction with HMGCR and Insig-1

The Insig proteins provide an ER-resident platform for sterol-dependent interactions between HMGCR and its regulatory components (Dong et al., 2012). RNF145 is sterol regulated and degrades HMGCR, making it important to determine whether it interacts directly with HMGCR, or via the Insig proteins. In sterol-replete but not sterol-deplete conditions, endogenous HMGCR co-immunoprecipitates both epitope-tagged RNF145 (**Figure 6A**, **Figure 3 – figure supplement 4C** lane 3 for relative RNF145-V5 levels upon reconstitution), as well as endogenous RNF145 (**Figure 6B**).

**Figure 6.**
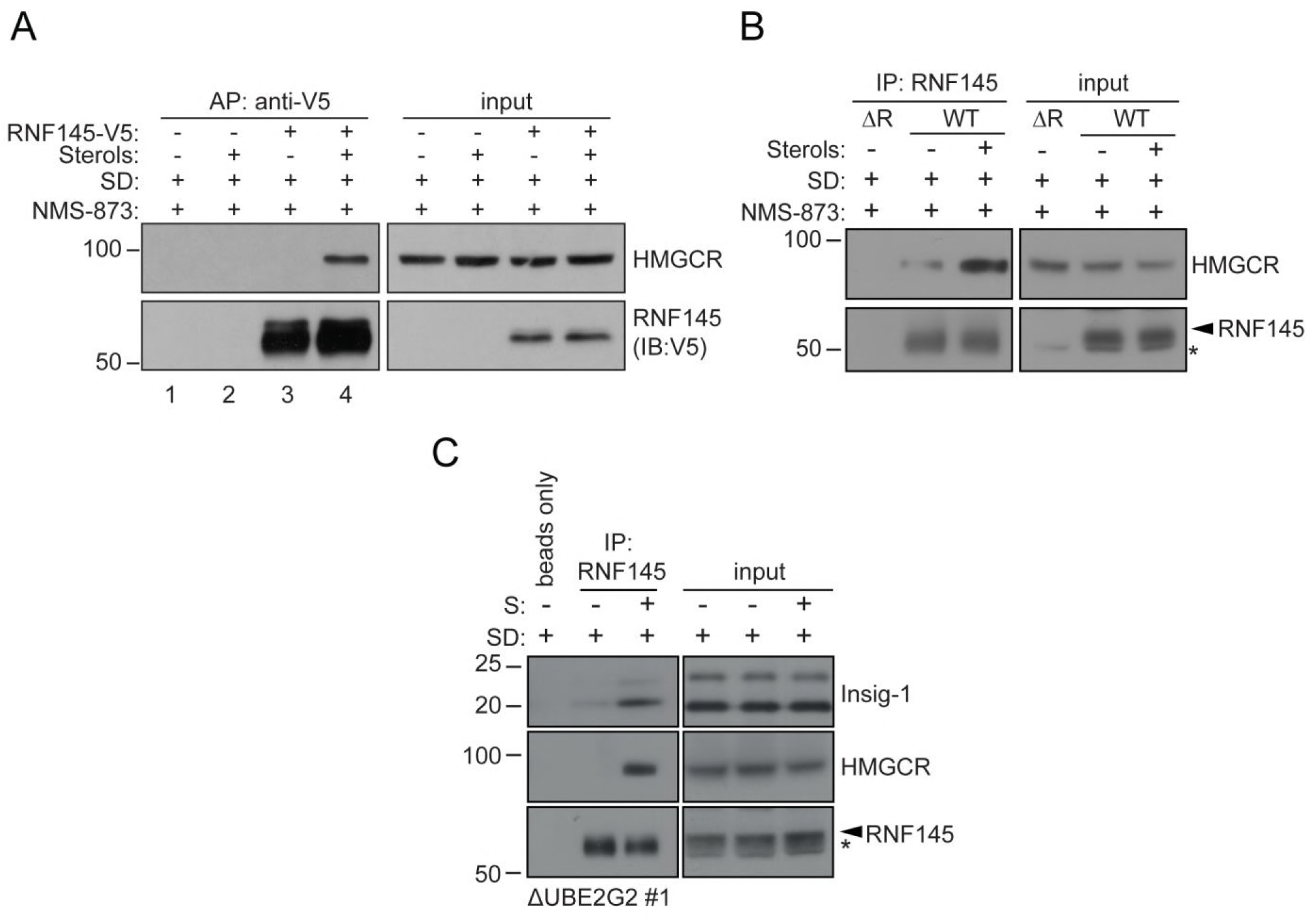
Endogenous RNF145 shows sterol-sensitive binding to Insig-1 and HMGCR. **(A)** Exogenous RNF145 shows sterol-sensitive binding to HMGCR. RNF145 knockout cells stably reconstituted with RNF145-V5 (ΔR145 #4 + R145-V5, as shown in **Figure 3 – figure supplement 4 C**, lane 3) were sterol-depleted (SD, 20h) and sterols (S) added back for 1h in the presence of NMS-873 (10 μM, 1.5h). RNF145-V5 was affinity-purified (AP) and HMGCR detected by immunoblotting. Representative of ≥ 3 independent experiments. **(B - C)** Endogenous RNF145 shows sterol-sensitive binding to HMGCR and Insig-1. **(B)** HeLa WT or ΔRNF145 #4 (ΔR) cells were treated as in **(A)** and endogenous RNF145 was immunoprecipitated (IP), and RNF145 and HMGCR detected by immunoblot analysis. Nonspecific bands are designated by an asterisk (*). Representative of ≥ 3 independent experiments. **(C)** HeLa UBE2G2 knockout cells (ΔUBE2G2 #1) were sterol-depleted (SD, 20h) and sterols (S) added for 1h. Endogenous RNF145 was affinity-purified and following SDS-PAGE separation, Insig-1 and HMGCR detected by immunoblot analysis. Representative of ≥ 2 independent experiments.

The low expression levels of endogenous RNF145 made any interaction with endogenous Insig-1 challenging to detect. We circumvented this problem by repeating the co-immunoprecipitation in UBE2G2 knockout cells, which express increased levels of endogenous RNF145 (**Figure 5B**). Under these conditions, RNF145 showed a sterol-dependent interaction with Insig-1, correlating with RNF145’s association with HMGCR (**Figure 6C**). Importantly, endogenous RNF145 is not, therefore, continually bound to Insig-1, but, like HMGCR, associates with Insig-1 in a sterol-dependent manner.

### In the absence of RNF145 and gp78, Hrd1 targets HMGCR for degradation

Despite our two genetic screens identifying a requirement for RNF145 and gp78 in HMGCR degradation (**Figure 2C** and **3B**), the combined loss of these two ligases failed to completely inhibit sterol-induced HMGCR degradation (**Figure 3C (iv)**; **Figure 7A (ii)**). Furthermore, ablation of UBE2G2 in RNF145/gp78 double-knockout cells further exacerbated the sterol-dependent degradation defect (**Figure 7A (iv)**), predicting the role for an additional E3 ligase(s) utilising UBE2G2 in HMGCR degradation. We therefore assessed whether ablation of either of the two remaining ER-resident E3 ligases that use UBE2G2, TRC8 (van de Weijer et al., 2017) and Hrd1 (Kikkert et al., 2004), exacerbated the HMGCR-degradation defect in RNF145/gp78 double-knockout cells (**Figure 7**, **Figure 7 – figure supplement 1B** and **2B** for knockdown validation). While the loss of TRC8 had no effect on HMGCR-Clover expression, the loss of Hrd1 in RNF145/gp78 double-knockout cells increased steady-state HMGCR-Clover expression and caused a complete block in the sterol-accelerated degradation of HMGCR-Clover (**Figure 7B (ii)**, **Figure 7 – figure supplement 1A** for validation with individual independent sgRNAs). The additive effect of Hrd1 depletion on the sterol-induced turnover of endogenous HMGCR was independently confirmed by immunoblot analysis (**Figure 7C**, compare lanes 2, 4 and 6) and was observed as early as 60 minutes after sterol addition (**Figure 7 – figure supplement 1D**, compare lanes 7 and 9). Importantly, depletion of Hrd1, either alone or in combination with depletion of either gp78 or RNF145, did not affect HMGCR-Clover degradation (**Figure 7 – figure supplement 1C**). Moreover, TRC8 depletion affected neither steady-state HMGCR-Clover expression, nor sterol-induced HMGCR-Clover degradation (**Figure 7B (iii)**). Indeed, despite a functional TRC8 depletion (**Figure 7 – figure supplement 2B** for validation) (Stagg et al., 2009), we could detect no role for TRC8, depleted either alone or in combination with RNF145, in the sterol-induced degradation of HMGCR (**Figure 7 – figure supplement 2A**).

**Figure 7.**
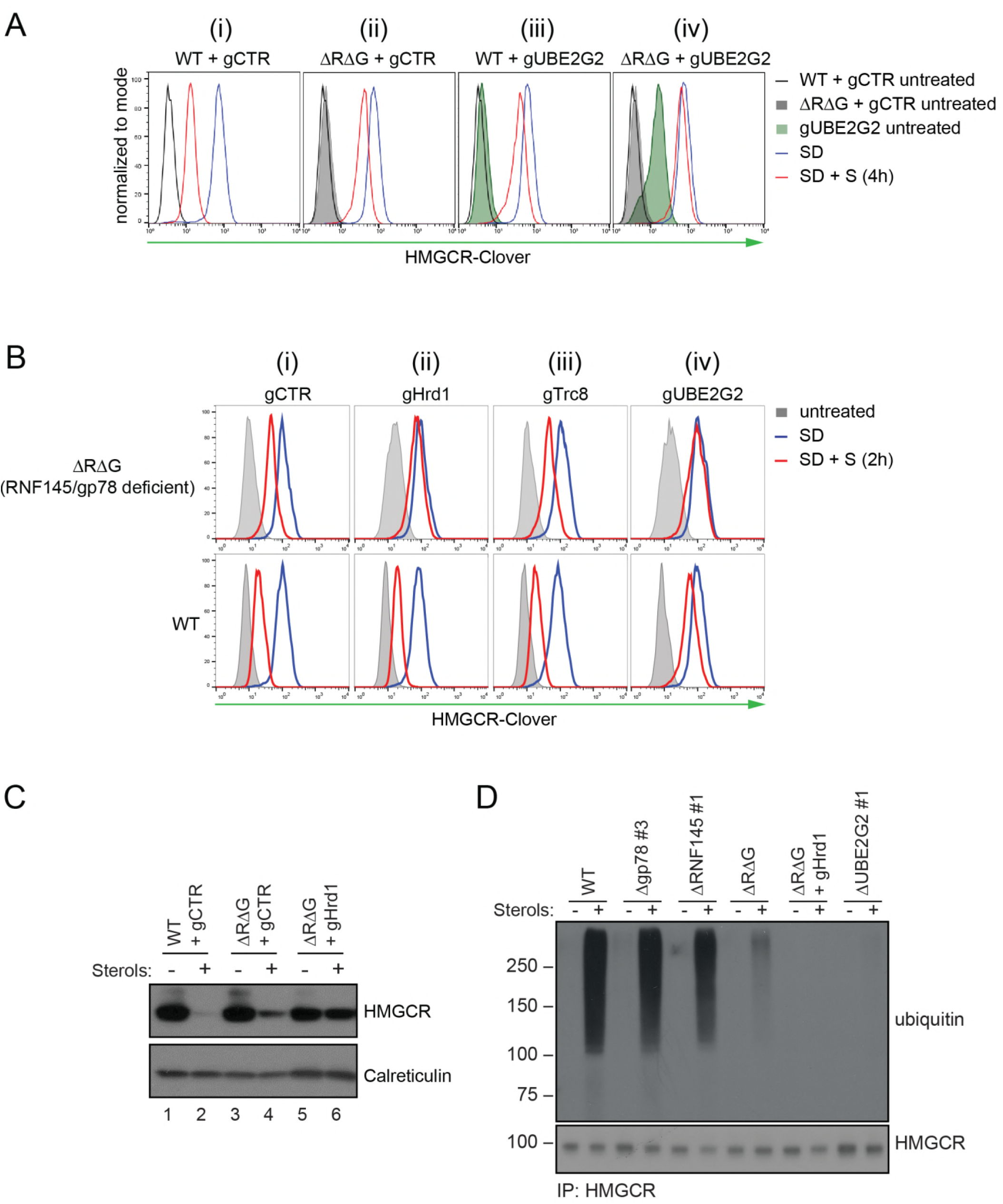
In the absence of RNF145 and gp78, Hrd1 targets HMGCR for ubiquitination and degradation. **(A)** Loss of gp78, RNF145 and UBE2G2 exert an additive effect on HMGCR degradation. WT or RNF145/gp78 double knockout (ΔRΔG #11) HeLa HMGCR-Clover cells transiently expressing gRNAs targeting UBE2G2 (gUBE2G2) or B2M (gCTR) were enriched by puromycin selection, sterol-depleted (SD) overnight and HMGCR-Clover expression assessed ± sterols (S, 4h). Representative of 3 independent experiments. **(B and C)** A targeted gene approach shows loss of Hrd1 from RNF145/gp78 double knockout cells blocks sterol-accelerated degradation of HMGCR. **(B)** WT and ΔRNF145 Δgp78 (ΔRΔG) HMGCR-Clover cells transfected with gCTR, a pool of four sgRNAs targeting either Hrd1 (gHrd1), or TRC8 (gTRC8) or gUBE2G2, were sterol-depleted (SD, 20h) and HMGCR-Clover expression assessed ± sterols (S, 2h). **(C)** WT and RNF145+gp78 double knockout cells (ΔRΔG #7) HMGCR cells were transfected with a pool of four Hrd1-specific sgRNAs or gCTR, and sterol-depleted overnight before addition of sterols (4h) and analysis by SDS-PAGE and immunoblotting. RNF145 and gp78 knockout validation is shown in **Figures 3 – figure supplement 1B** (ΔRNF145#1) and **Figure 3 – figure supplement 3B** (ΔRΔG#7), respectively. **(D)** RNF145, gp78 and Hrd1 are required for sterol-accelerated HMGCR ubiquitination. HMGCR was immunoprecipitated (IP) from the indicated cell lines grown in sterol-depleted media (20h) ± sterols (2h). MG-132 (50 μM) was added 30 minutes before sterol supplementation. Ubiquitinated HMGCR was detected using an anti-ubiquitin antibody.

In summary, gp78 with RNF145 are the only combination of ligases whose loss inhibited HMGCR degradation. Hrd1 depletion also delays sterol-induced HMGCR degradation, but only in the absence of RNF145 and gp78.

### RNF145, gp78 and Hrd1 are required for sterol-accelerated HMGCR ubiquitination

As a complete block of sterol-accelerated HMGCR degradation required the depletion of all three UBE2G2-dependent ligases, we determined how the sequential depletion of these ligases affected the ubiquitination status of HMGCR. The combined loss of RNF145 with gp78 showed a dramatic reduction in HMGCR ubiquitination, but a complete loss of ubiquitination required the depletion of all three ligases (**Figure 7D**). As predicted, depletion of UBE2G2 also caused a marked decrease in HMGCR ubiquitination. Taken together, these results demonstrate the remarkable plasticity of the HMGCR-degradation machinery.

## DISCUSSION

The generation of a dynamic, cholesterol-sensitive endogenous HMGCR reporter cell line allowed an unbiased genetic approach to identify the cellular machinery required for sterol-accelerated HMGCR degradation. This reporter cell line has the advantage of being able to identify both complete and partial phenotypes and helps explain why the identity of the E3 ligases responsible for the sterol-accelerated degradation of HMGCR has remained controversial. We find that three E3 ligases - RNF145, gp78 and Hrd1 - are together responsible for HMGCR degradation (**Figure 8**). The activity of the two primary ligases, RNF145 and gp78 is partially redundant as the loss of gp78 alone did not affect HMGCR degradation, while loss of RNF145 showed only a small reduction on HMGCR degradation. In the absence of both RNF145 and gp78, a third ligase, Hrd1, can compensate and partially regulate HGMCR degradation, but this effect of Hrd1 is only revealed in the absence of both RNF145 and gp78, and in no other identified combination.

**Figure 8.**
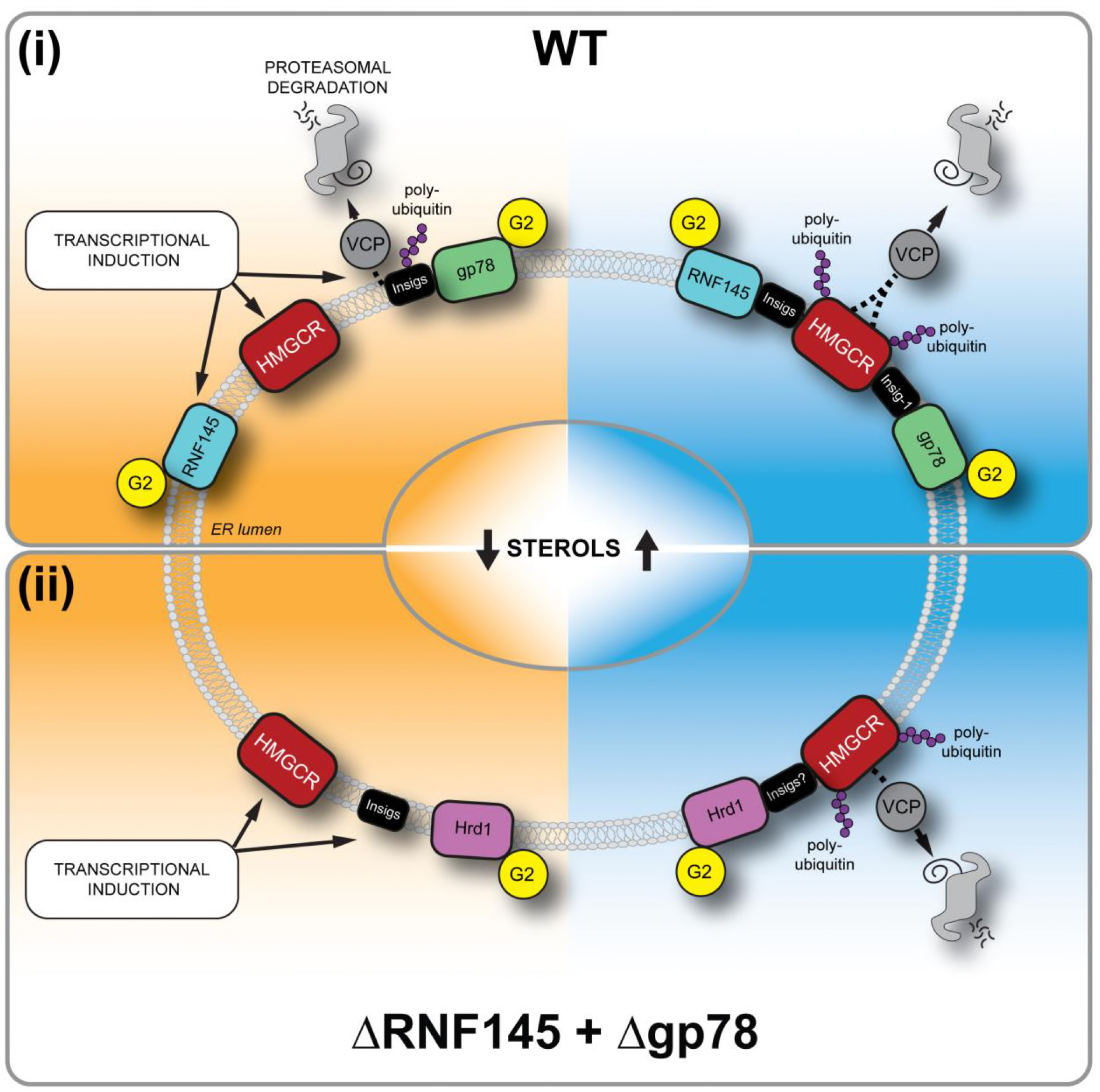
Sterol-induced degradation by RNF145, gp78, and Hrd1. (i) Under sterol-depleted conditions (shaded orange), HMGCR, Insig1, and RNF145 are transcriptionally induced leading to accumulation of RNF145 and HMGCR. Insigs are continually turned over by gp78-mediated polyubiquitination, extracted from the membrane by VCP and degraded by the 26S proteasome. HMGCR stability is dramatically increased as it is not engaged by either RNF145, gp78 or their shared E2 ubiquitin ligase UBE2G2 (G2). In the presence of sterols (shaded blue), RNF145 and gp78 are recruited to HMGCR in an Insig-assisted fashion, mediating the sterol-accelerated and UBE2G2-dependent degradation of HMGCR by the UPS. Under these conditions both RNF145 and gp78 can independently ubiquitinate HMGCR, which is then extracted from the ER membrane in a VCP-dependent manner. The stoichiometry and make-up of the different Insig complexes within the ER membrane are unknown (ii) When RNF145 and gp78 are not available, Hrd1 and UBE2G2 can promote removal of HMGCR in the presence of sterols.

Initial reports of a role for gp78 in HMGCR degradation, either alone (Song, Sever, & DeBose-Boyd, 2005) or in combination with TRC8 (Jo et al., 2011), were not reproduced in an independent study (Tsai et al., 2012) and so this important issue has remained unresolved. Our initial genome-wide screen successfully identified many of the components known to be required for sterol-accelerated HMGCR degradation (e.g. Insig-1/2, and AUP1, **Figure 2C**) (Sever et al., 2003b; Miao et al., 2010; Jo et al., 2013), thus validating the suitability of this genetic approach. The screen also identified the E2 conjugating enzyme UBE2G2 and the E3 ligase RNF145. Depletion of UBE2G2 prevented HMGCR degradation, implying that all ligases involved in HMGCR degradation utilise this E2 enzyme. In contrast, and despite being a high confidence hit in our screen, depletion of RNF145 caused a highly reproducible but small inhibition of sterol-accelerated degradation, confirming the sensitivity of the screen to detect partial phenotypes and predicting the requirement for at least one additional UBE2G2-dependent ligase. A subsequent, targeted ubiquitome library screen in an RNF145-knockout reporter cell line confirmed a role for gp78 in HMGCR degradation. Gp78 has previously been shown to use UBE2G2 as its cognate E2 enzyme in the degradation of ERAD substrates (Chen et al., 2006). During preparation of this manuscript, the combined involvement of RNF145 and gp78 in HMGCR degradation in hamster (CHO) cells was also reported (Jiang et al., 2018), confirming the role for these ligases in other cell lines.

The availability of an RNF145-specific polyclonal antibody provides further insight into the expression and activity of endogenous RNF145, without the concerns of overexpression artefacts. RNF145 is an ER-resident E3 ligase with several unique features that make it well-suited for HMGCR regulation. A challenge facing all proteins responsible for cholesterol regulation is that the target they monitor, cholesterol, resides entirely within membranes. We find that RNF145, like SCAP and HMGCR, two key proteins involved in cholesterol regulation, is both sterol regulated and associates with its Insig binding partner in a sterol-responsive manner. Under sterol deplete conditions, endogenous RNF145 is associated with neither HMGCR nor Insig1, but the addition of sterols triggers RNF145 binding to the ER-resident Insig-1 protein (**Figure 5C**). Like HMGCR and SCAP, RNF145 contains a sterol-sensing domain in its transmembrane region (Cook et al., 2017), suggesting that sterols facilitate its association with Insigs. Similarly, the association of HMGCR with Insigs and gp78 is also sterol-dependent through its SSD (Lee et al., 2007). Therefore, sterols trigger the recruitment of RNF145 to HMGCR, leading to Insig-dependent HMGCR ubiquitination and degradation. This ability of RNF145 to rapidly bind Insigs following sterol availability is a feature shared with the related sterol-responsive proteins including HMGCR and SCAP and further supports a key role for this ligase in HMGCR regulation.

A striking feature of RNF145 is its short half-life and rapid proteasome-mediated degradation, which contrasts with the long-lived gp78 (**Figure 6A**, **Figure 5 – figure supplement 1B**). RNF145 is an intrinsically unstable ligase whose half-life is regulated through autoubiquitination and was not prolonged on binding to Insig proteins (data not shown). Its stability and turnover is RING- and UBE2G2-dependent, but independent of either the gp78 (**Figure 6A-C**) or Hrd1 ligase (**Figure 5 – figure supplement 1C**). As cells become sterol-depleted, the transcriptional increase in RNF145 (**Figure 6E**) likely anticipates the need to rapidly eliminate HMGCR, once normal cellular sterol levels are restored.

While it is not unusual for more than one ligase to be required for substrate ERAD degradation (Christianson and Ye, 2014; Morito et al., 2008; Stefanovic-Barrett et al., 2018), the redundancy in HMGCR turnover is intriguing. This may simply reflect the central role of HMGCR in the mevalonate pathway and the importance of a fail-safe mechanism of HMGCR regulation to both maintain substrates for non-sterol isoprenoid synthesis and prevent cholesterol overproduction. Alternative explanations can also be considered, particularly as the properties of RNF145 and gp78 are so different. Under sterol-deplete conditions gp78 also regulates the degradation of Insig-1, but following addition of sterols, the association of Insigs with SCAP displaces its binding to gp78 (Yang et al., 2002; Lee et al., 2006). Different Insig-associated complexes are therefore likely to co-exist within the ER membrane, under both sterol-replete and -deplete conditions, and therefore reflect the sterol microenvironment of the ER (Goldstein et al., 2006). Under these circumstances it might be advantageous to have more than one ligase regulating HMGCR. Alternatively, gp78 may provide basal control of the reductase, which can then be ‘fine-tuned’ by the sterol-responsive RNF145, reflecting the sterol concentration of the local ER environment. It will therefore be important to further understand the stoichiometry and nature of the different Insig complexes within the ER membrane. While all cells need to regulate their intracellular cholesterol, the contribution of each ligase to sterol regulation may also depend on their differential tissue expression. In this regard, liver-specific ablation of gp78 in mice has been reported to lead to increased steady-state levels of hepatocyte HMGCR (Liu et al., 2012), whereas, gp78 knockout MEFs show no apparent impairment in HMGCR degradation (Tsai et al., 2012). Further delineation of the contribution of each ligase to HMGCR degradation in different tissues and cell types will be important.

A role for the Hrd1 E3 ligase in HMGCR regulation was unanticipated, and both orthologues (gp78 and Hrd1) of yeast Hrd1p, which regulates yeast HMGCR (Hmg2p), are therefore involved in mammalian HMGCR turnover. The best recognised function of Hrd1 is the ubiquitination of misfolded or unassembled ER-lumenal and membrane proteins targeted for ERAD (Sato et al., 2009; Tyler et al., 2012; Christianson et al., 2008). Our finding that Hrd1 is only involved in HMGCR regulation when the other two ligases are absent, suggests that under sterol-rich conditions, and in the absence of RNF145 or gp78, conformational changes in the sterol sensing domains of HMGCR to a less ordered state are recognised and targeted by the Hrd1 quality control pathway.

In summary, our unbiased approach to identify proteins involved in sterol-regulated HMGCR degradation resolves the ambiguity of the E3 ligases responsible, and further unveils additional control points in modulating the activity of this important enzyme in health and disease.

## AUTHOR CONTRIBUTIONS

PJL, SAM, NV and RTT conceived the project. Experiments were carried out by SAM, NV, DJB and ASD. The CRISPR/Cas9 ubiquitin library was designed by JAN and SAM and generated by JAN and DJB. SAM, NV and PJL analysed the data. SAM and NV prepared the figures. NV, PJL and SM wrote the manuscript.

## ACKNOWLEDGEMENTS

We are grateful to the following for their help in this study: Michael Bassik (Stanford University) kindly shared the genome-wide CRISPR/Cas9 sgRNA library. Ron Kopito (Stanford University) kindly donated the pDonor loxP Ub-Puro plasmid. FACS experiments were enabled by R. Schulte and his FACS core facility team in CIMR. Stuart Bloor (CIMR), Gordon Dougan, Richard Rance and Nathalie Smerdon (Sanger Institute) assisted with Illumina sequencing. This work was supported by the Wellcome Trust, through a Principal Research Fellowship to P.J.L (210688/Z/18/Z), a Wellcome Trust Senior Clinical Research Fellowship to J.A.N. (102770/Z/13/Z) and a Welcome Trust PhD studentship to S.M.. The CIMR is in receipt of a Wellcome Trust strategic award (100140).

## ABBREVIATIONS

AMFR: - Autocrine Motility Factor Receptor
B2M: - beta-2-microglobulin
Chol: - cholesterol
CHX: - cycloheximide
CRISPR: - clustered regularly interspaced short palindromic repeats
CTR: - control
ER: - endoplasmic reticulum
ERAD: - ER-associated degradation
FCS: - fetal calf serum
Gp78: - glycoprotein 78
HMGCR: - 3-hydroxy-3-methyl-glutaryl-coenzyme A reductase
Hrd1: - HMG-CoA Reductase Degradation 1
IAA: - iodoacetamide
Insig-1/2: - Insulin-induced gene-1/2
LPDS: - lipoprotein-deficient serum
MBCD: - methyl-β-cyclodextrin
RNF145: - RING finger protein 145
SD: - sterol-depleted
SREBP2: - Sterol Regulatory Element Binding transcription factor 2
TRC8: - Translocation in renal carcinoma on chromosome 8
UBE2G2: - Ubiquitin-conjugating enzyme E2 G2
UPS: - ubiquitin proteasome system
VCP: - Valosin-containing protein
WT: - wild-type

---

**Figure 2 - figure supplement 1.**
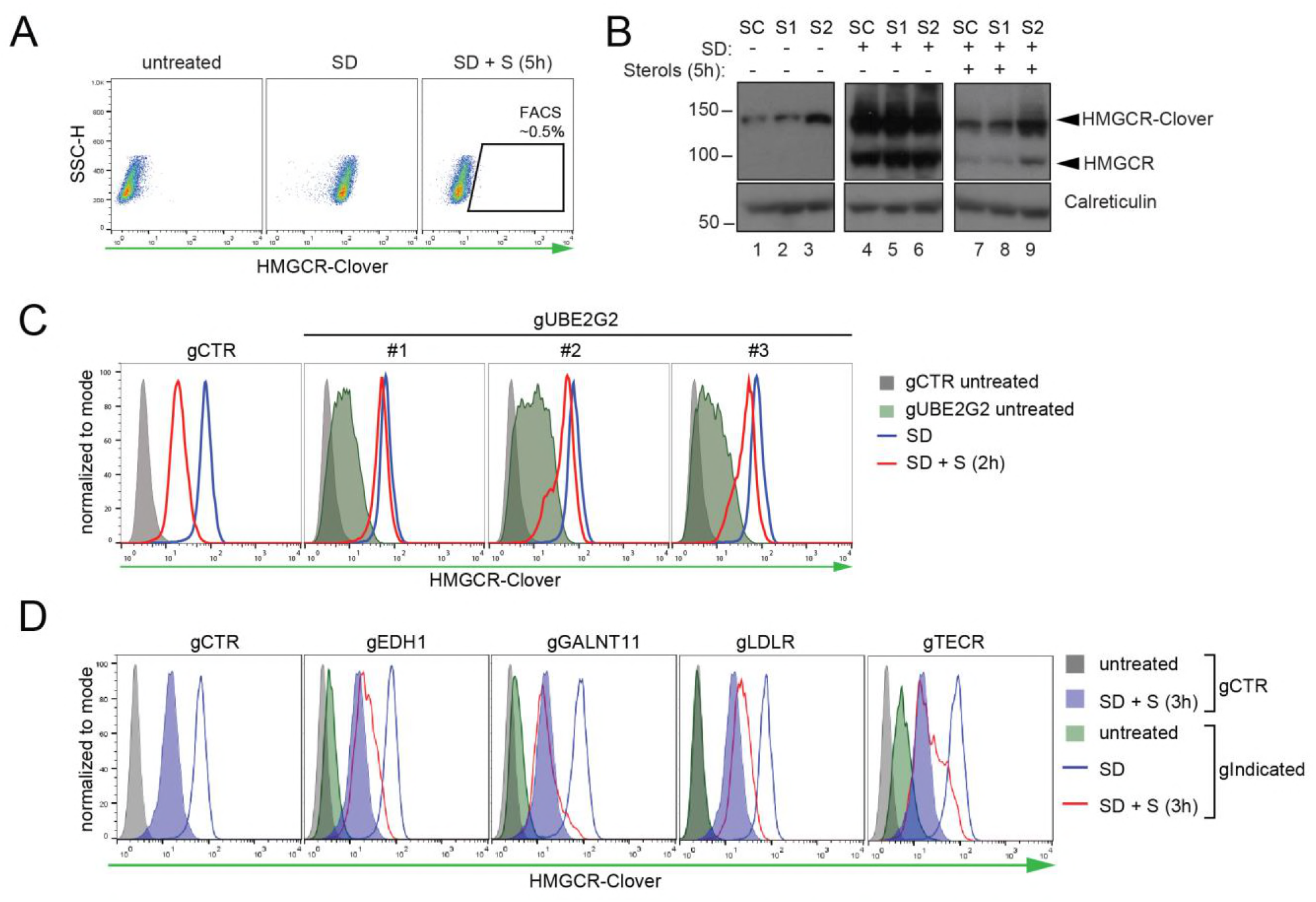
Genome-wide screen for proteins involved in HMGCR ERAD. **(A)** Gating strategy to enrich for Clover^high^ mutants with impaired sterol-induced HMGCR-Clover degradation. HeLa HMGCR-Clover cells mutagenized with sgRNA library were subjected to overnight sterol depletion (SD) before adding back sterols (SD + S) for 5h. Typically, the highest ~ 0.5% of Clover^high^ cells were selected for enrichment (indicated). **(B)** Immunoblot analysis for HMGCR-Clover enrichment after sort 1 (S1), sort (S2) as compared to the starting clone (SC). Cells were sterol-depleted (SD) overnight ± sterols (2h). **(C)** Flow cytofluorometric analysis of HMGCR-Clover cells transiently expressing three independent sgRNAs (gUBE2G2 #1-3) *versus* gB2M (gCTR) after overnight sterol-depletion ± sterols (2h). **(D)** HMGCR-Clover cells were transfected with a pool of four sgRNAs for each indicated gene or a sgRNA against B2M (gCTR) and treated as in (C).

**Figure 2 - figure supplement 2.**
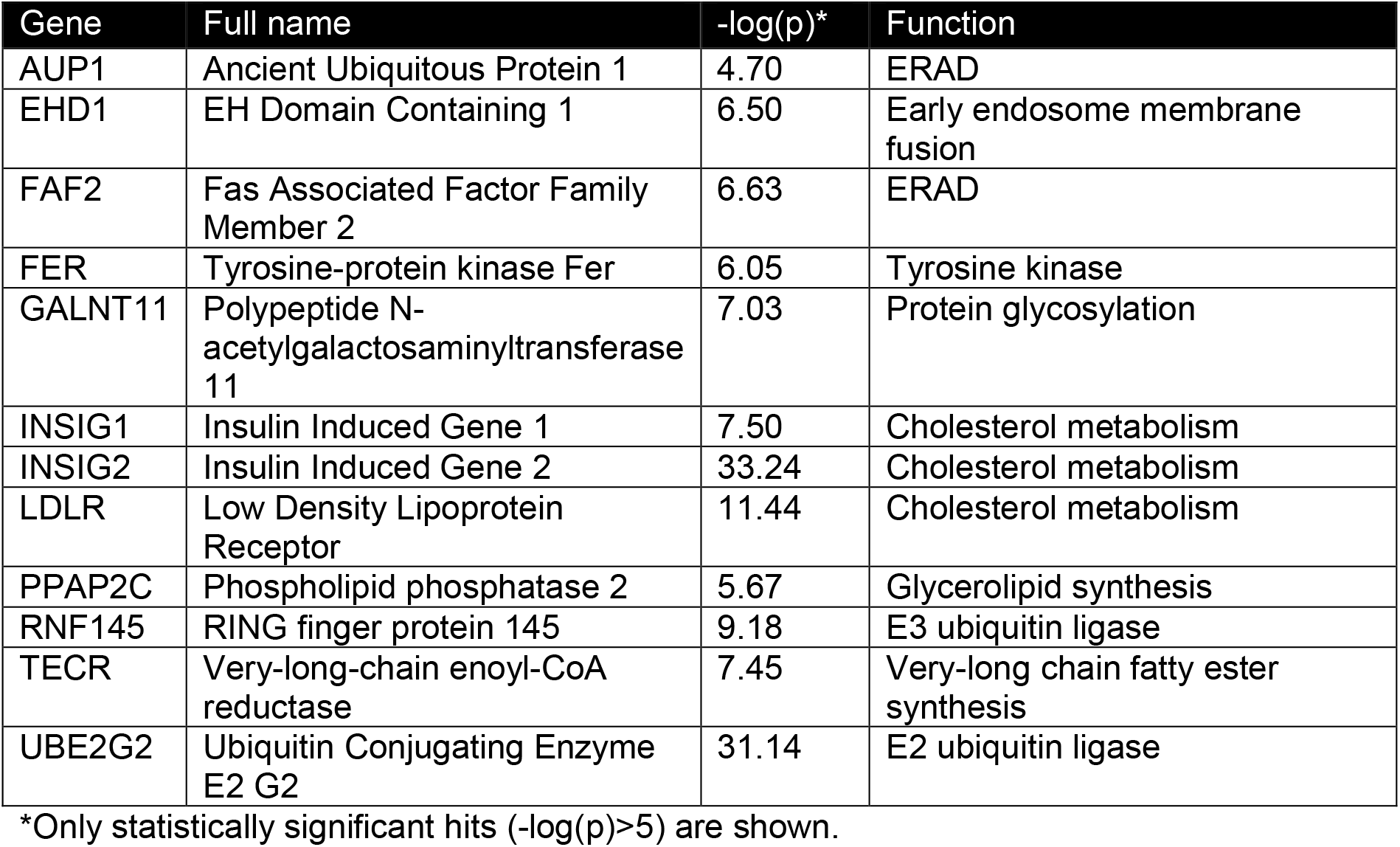
Candidate genes (- log(p) ≥ 5) identified in a genome-wide CRISPR/Cas9 screen for proteins involved HMGCR degradation.

**Figure 3 - figure supplement 1.**
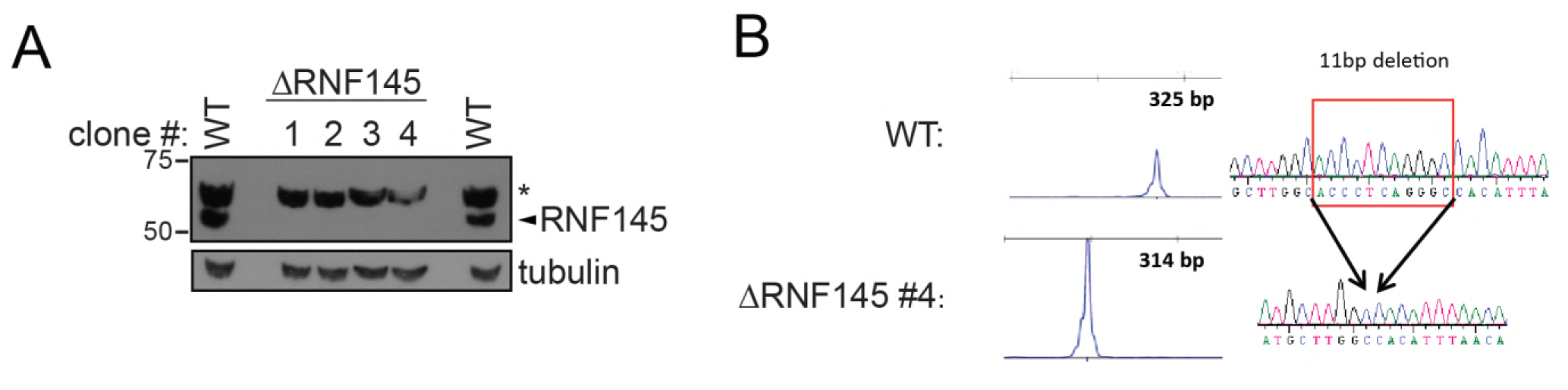
Validation of RNF145 knockout clones. **(A)** Immunoblot for RNF145 in WT *vs*. four RNF145 HeLa knockout (ΔRNF145) clones, generated with 2 independent sgRNAs. Non-specific bands are indicated (*). **(B)** Confirmation of RNF145 knockout clone #4 (ΔRNF145 #4). The genomic region surrounding the predicted sgRNA annealing site was amplified using fluorescent primers and amplicon size was determined capillary electrophoresis (Agilent Bioanalyzer 2100). Fluorescent traces are shown alongside amplicon sequences as obtained by Sanger sequencing, confirming an 11 bp deletion (indicated).

**Figure 3 - figure supplement 2.**
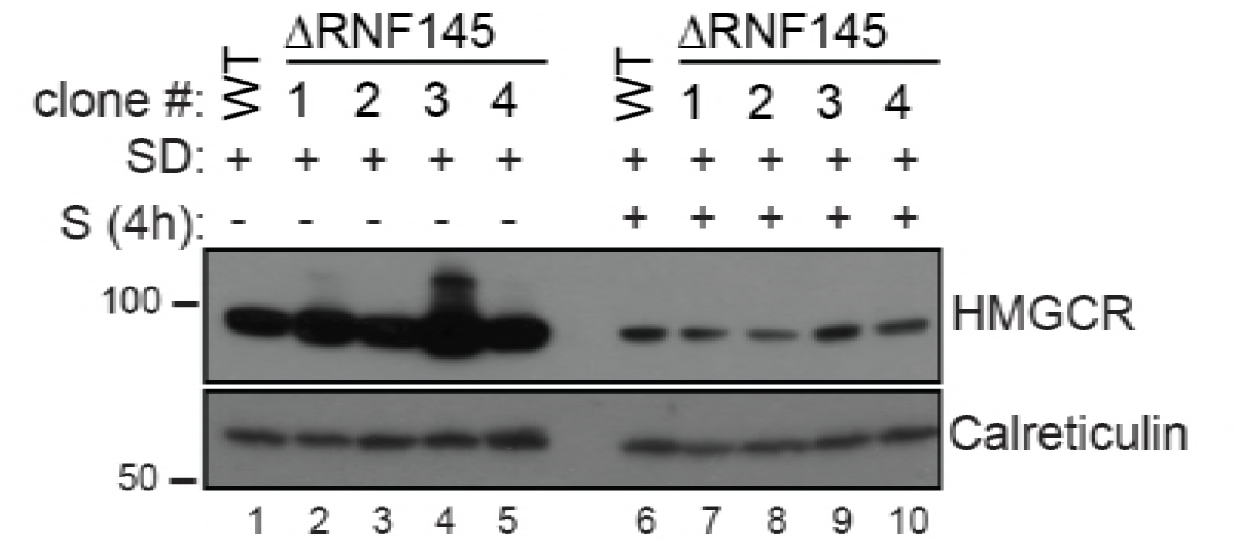
RNF145 loss is insufficient to block sterol-induced HMGCR degradation. Cells were sterol-depleted (SD) overnight before addition of sterols (S, 4h). Whole-cell lysates from WT and four RNF145 knockout clones (#1-4) were separated by SDS-PAGE and HMGCR levels visualised by immunoblot analysis. Calreticulin serves as a loading control.

**Figure 3 - figure supplement 3.**
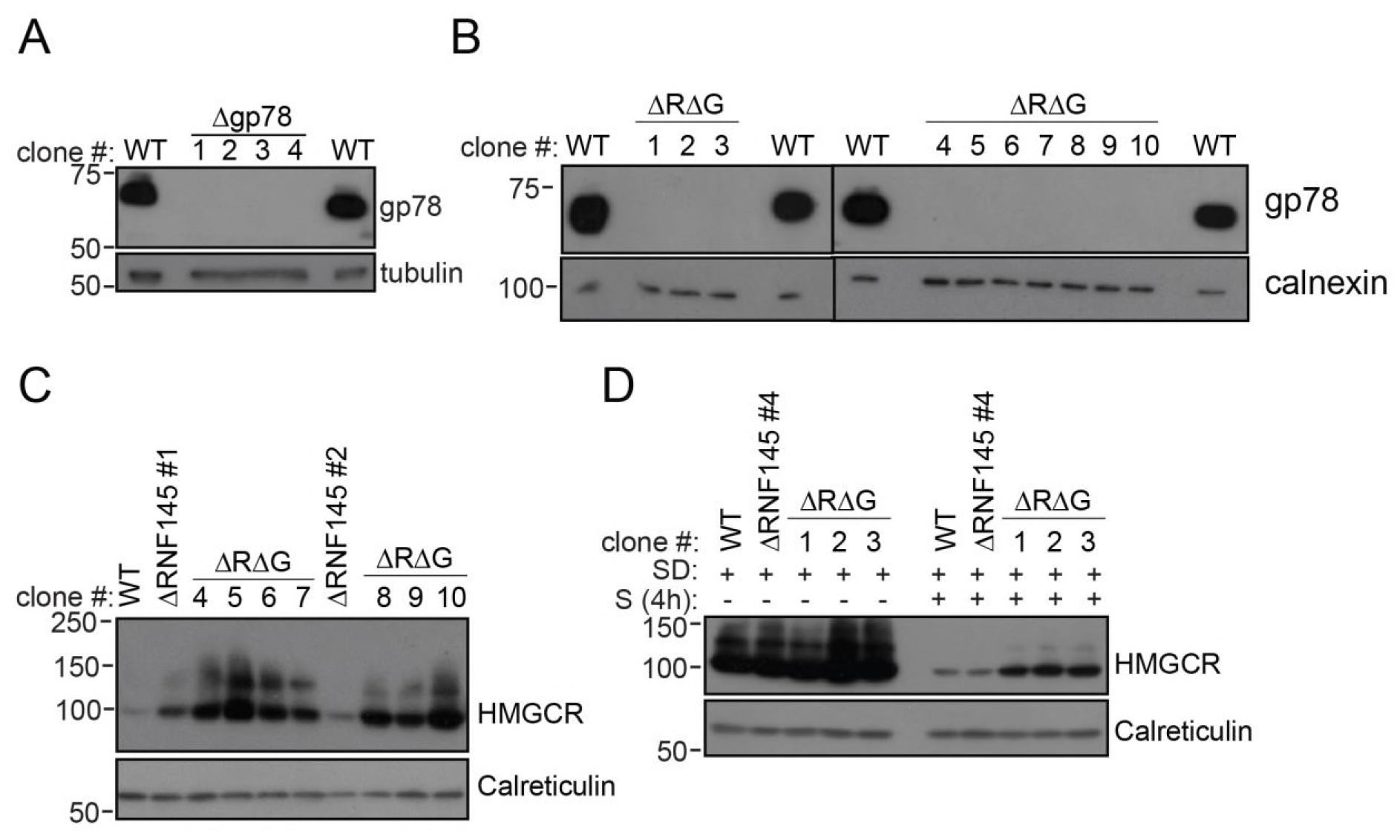
RNF145/gp78 double-knockout cells show increased HMGCR at steady-state and impaired sterol-induced HMGCR degradation. **(A)** Immunoblot of four gp78 knockout HeLa clones derived by transfection with two independent gp78 sgRNAs. **(B)** Confirmation of gp78 loss in RNF145/gp78 knockout HeLa cells derived from ΔRNF145 clones #1, #2, and #4 (for validation of RNF145 knockout see **Figure 3 – figure supplement 1**). Calnexin serves as a loading control. **(C)** Steady-state expression of HMGCR in HeLa WT, ΔRNF145, and RNF145/gp78 double knockout clones (ΔRΔG) was determined by immunoblotting. **(D)** Indicated cell lines were sterol-depleted (SD) overnight ± sterols (S, 4h) and HMGCR detected by immunoblotting. Calreticulin serves as a loading control. LE, long exposure.

**Figure 3 - figure supplement 4.**
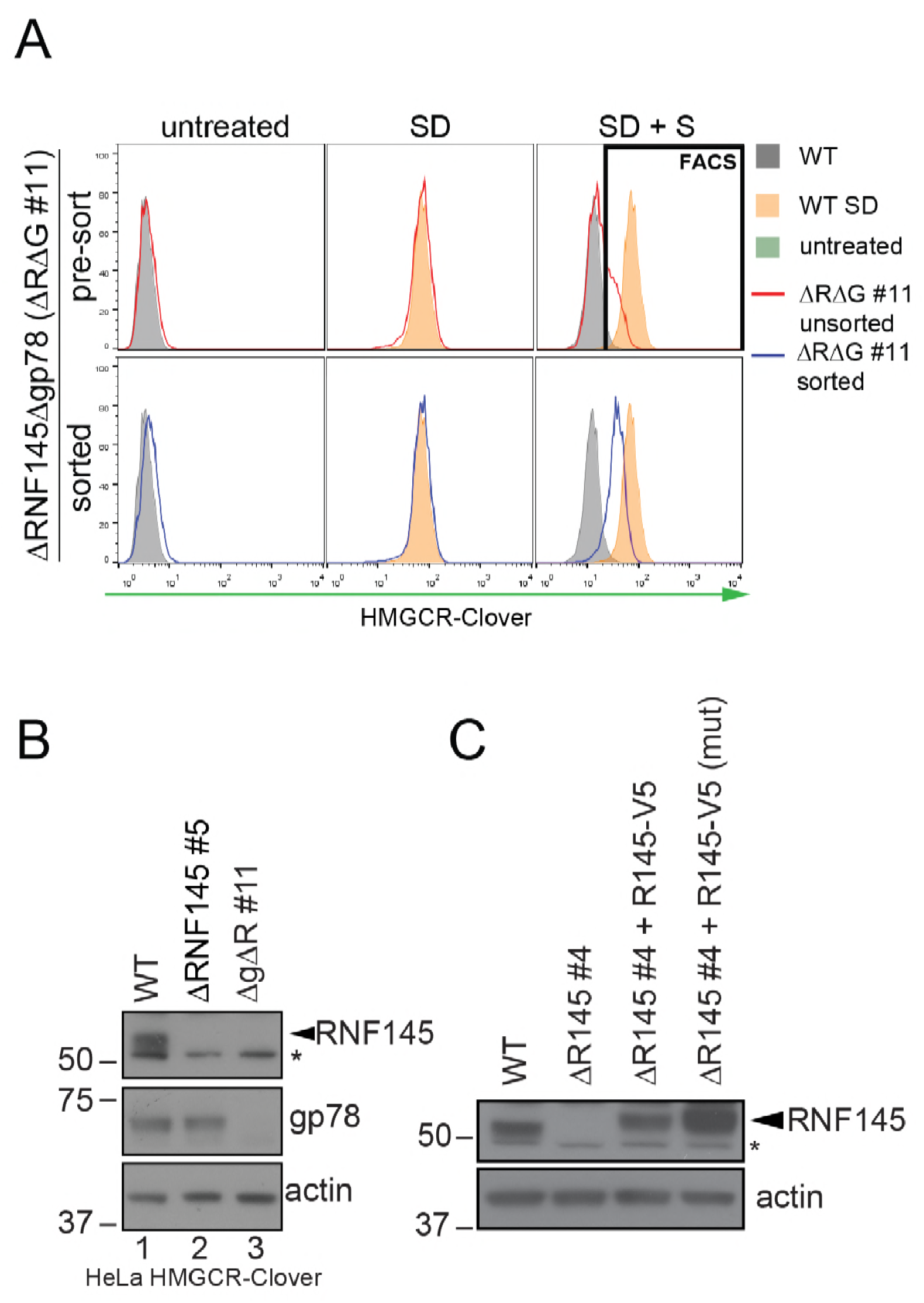
Establishment of RNF145/gp78 knockout HeLa HMCR-Clover and RNF145 complementation cell lines. **(A)** HMGCR-Clover cells were transfected with RNF145 sgRNA#8 and gp78 sgRNA#4 (ΔRΔG #11, for sgRNAs used see **Supplementary Files 4 and 5**) and knockout pools were enriched with puromycin. Eight days *post* transfection, cells were sterol-depleted (SD) overnight ± sterols (S, 2h). HMGCR-Clover^high^ cells were enriched by FACS. **(B)** RNF145 and gp78 levels in WT *versus* ΔRNF145 clones #5, and RNF145/gp78 depleted (ΔRΔG #11) HeLa HMGCR-Clover cells. **(C)** Stable genetic complementation of ΔRNF145 (clone #4) cells by transduction with constructs encoding either RNF145-V5 (ΔR145 #4 + R145-V5) or RNF145(C552A, H554A)-V5 (ΔR145 #4 + R145-V5 (mut)). RNF145 variants were detected using an RNF145-specific antibody. Non-specific bands are indicated (*).

**Figure 3 - figure supplement 5.**
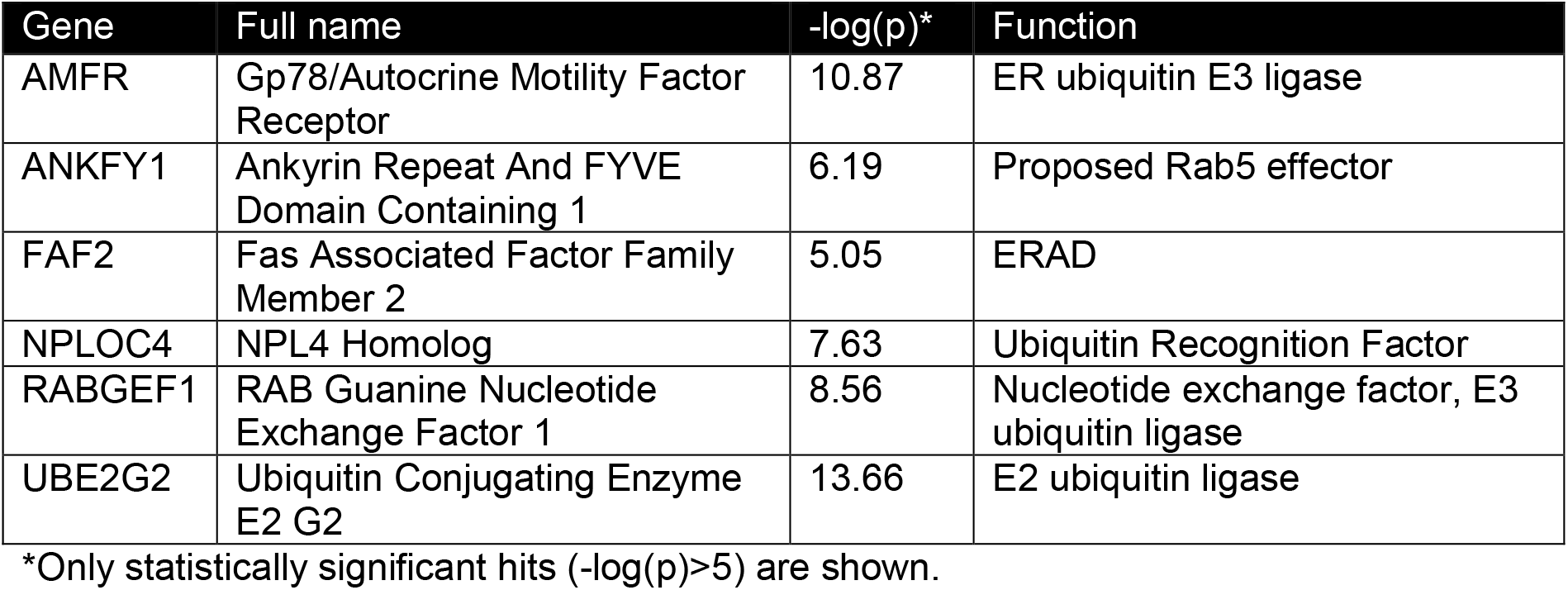
Candidate genes (-log(p)≥5) identified in a ubiquitome CRISPR/Cas9 screen for proteins mediating HMGCR degradation in RNF145 deficient cells.

**Figure 5 - figure supplement 1.**
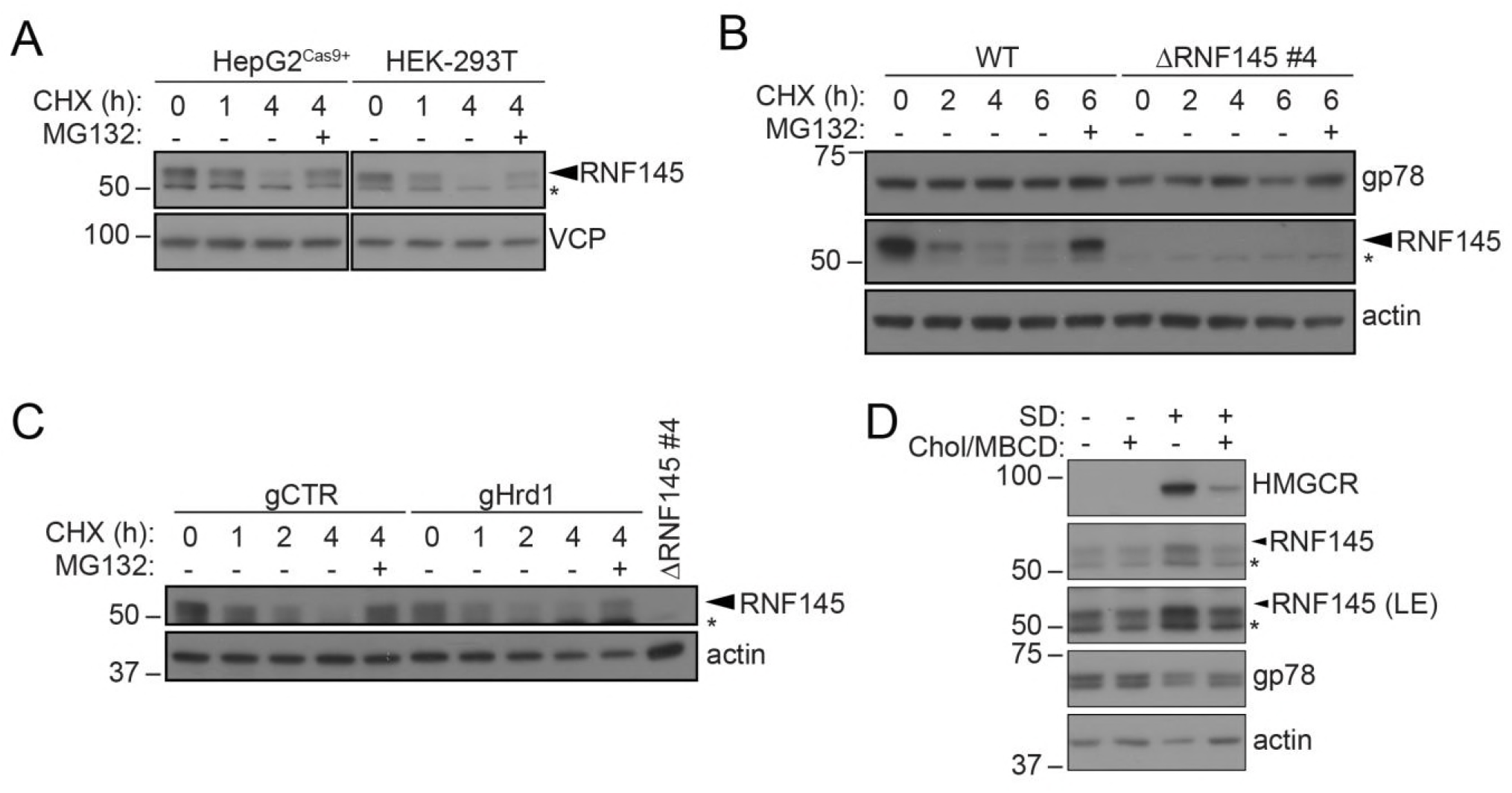
RNF145 is rapidly degraded by the ubiquitin proteasome system. **(A)** Translation shutoff assay in HepG2^Cas9+^ and HEK-293T cells. Cells were treated with cycloheximide (CHX, 1 μM) ± MG132 (20 μg/ml) for the indicated times and endogenous RNF145 levels determined by immunoblotting. **(B)** Gp78 is stable in the absence of RNF145. WT and ΔRNF145 #4 HeLa cells were cultured in the presence of CHX, (1 μM) ± MG132 (10 μg/ml) for 0-6 h and gp78/RNF145 levels monitored by Western blotting. The asterisk (*) indicates a non-specific band. **(C)** HeLa^Cas9^+ HMGCR-Clover were transiently transfected with a pool of 4 Hrd1-specific sgRNAs (gHrd1) or a B2M targeting control sgRNA (gCTR) and sgRNA containing cells enriched by puromycin selection. Cells were treated as in **(B)** for the indicated times. **(D)** HeLa cells were grown under sterol-rich or sterol-deplete conditions for 48 h in the presence of mevastatin (10 μM) and mevalonate (50 μM) ± complexed cholesterol (chol:MBCD, 25 μM). SDS-PAGE and immunoblot analysis was performed on whole-cell lysates. Non-specific bands are indicated (*). LE, long exposure.

**Figure 7 - figure supplement 1.**
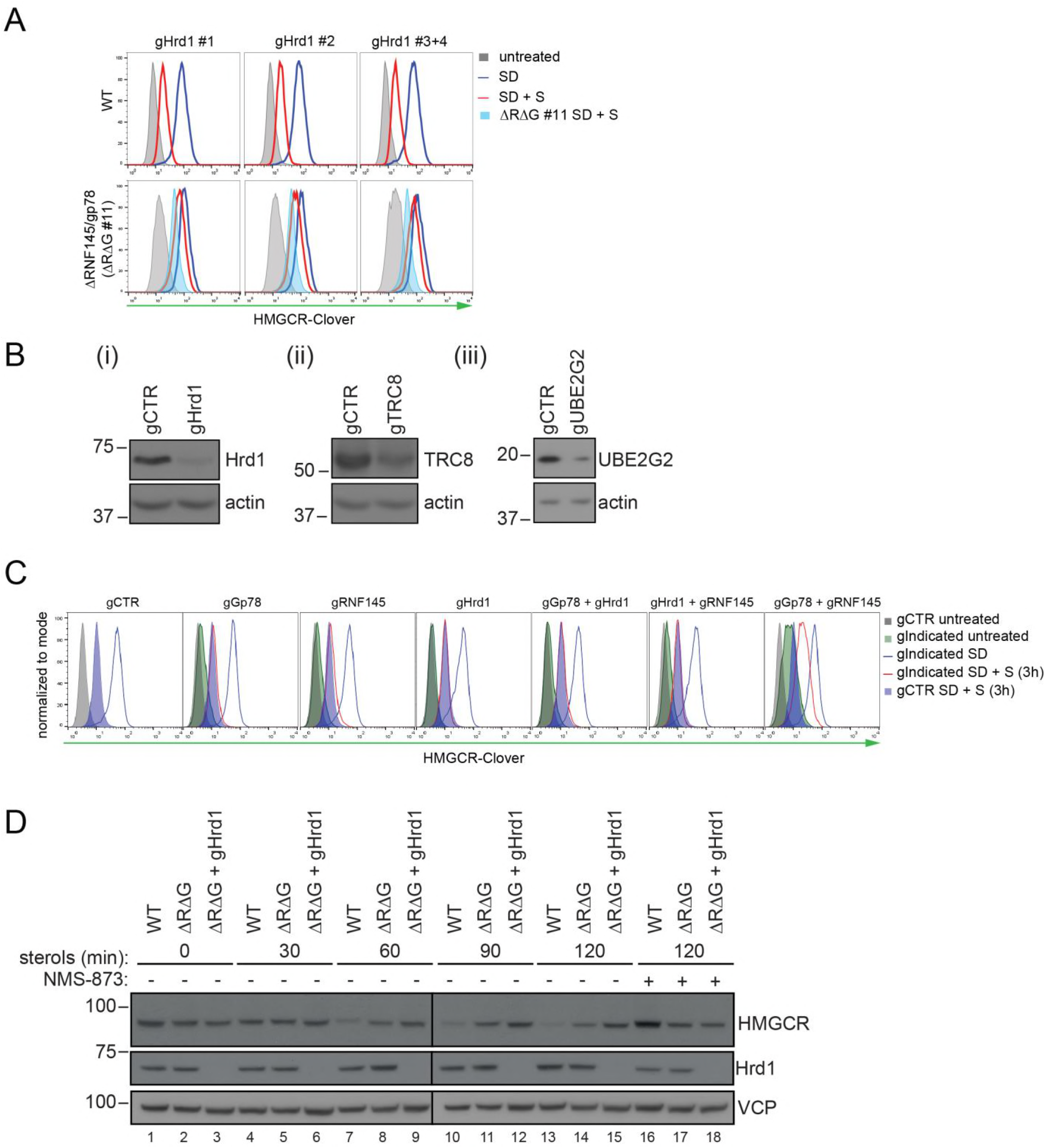
Combinatorial depletion of E3 ligases. **(A)** WT or RNF145/gp78 knockout (ΔRΔG #11) HeLa HMGCR-Clover cells transiently transfected with a B2M-specific (gCTR) or Hrd1-specific (gHrd1 #1-4) sgRNAs were sterol-starved (SD, 20h) ± sterols (S, 2h). HMGCR clover expression was detected by FACS analysis. Cells transfected with gCTR are from the same experiment shown in **Figure 7B** and histograms (gCTR SD+S) were therefore re-plotted. **(B)** Validation of Hrd1 (i), TRC8 (ii), and UBE2G2 (iii) depletion in ΔRΔG #11 cells. Cell lines used in **Figure 7B** and **Figure 7 – figure supplement 1A** were collected at steady-state and indicated proteins detected from whole-cell lysate by immunoblotting. A sgRNA targeting B2M (gCTR) served as a control. **(C)** WT HMGCR-Clover transfected with indicated guide pools were sterol-depleted (SD) overnight ± sterols (S, 3h). Reporter expression was measured by FACS. **(D)** WT and ΔRΔG#7 HeLa cells transfected with gCTR or a pool of four Hrd1-specific sgRNAs were sterol-depleted (20h) before addition of sterols for the indicated times ± NMS-873 (10 μM). Validation of RNF145 (ΔRNF145#1) and gp78 knockout (ΔRΔG#7) can be found in **Figure 3 – figure supplement 1A** and **3B**, respectively.

**Figure 7 - figure supplement 2.**
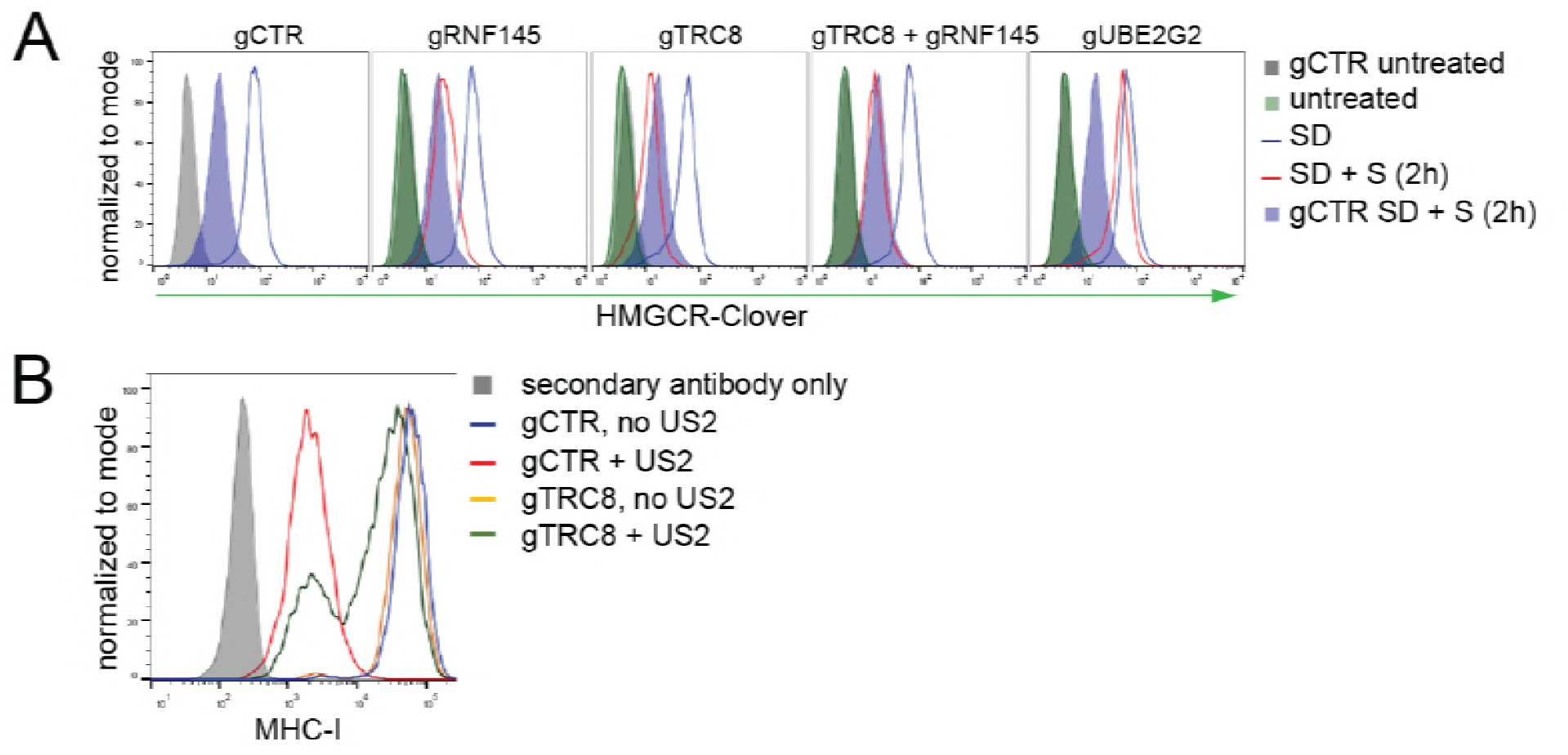
TRC8 depletion does not affect HMGCR-Clover degradation. **(A)** Overnight sterol depletion (SD) ± sterols (S, 2h) in HeLa HMGCR-Clover cells transiently transfected with pools of indicated guides as described in Materials and Methods. **(B)** TRC8 knockdown was confirmed by US2-mediated TRC8-dependent downregulation of MHC-I. HeLa cells transiently expressing either control sgRNA (gCTR) or gTRC8 were selected for puromycin resistance and transduced with a lentiviral US2 and/or TRC8 construct 5 days *post* transfection. Cell-surface MHC-I staining and FACS analysis were performed on day 10 *post* transfection.

## MATERIALS AND METHODS

### Plasmids and expression constructs

Single guide RNAs (sgRNAs) were cloned into pSpCas9(BB)-2A-Puro V1 (Addgene #48139, deposited by Dr. Feng Zhang), pSpCas9(BB)-2A-Puro V2 (Addgene #62988, deposited by Dr. Feng Zhang) as previously described (Ran et al., 2013). To generate the ubiquitome sgRNA library, sgRNAs (sgRNA sequences in **Supplementary File 1**) were cloned into pKLV-U6gRNA(BbsI)-PGKpuro2ABFP (Addgene # 50946) as reported previously (Doench et al., 2016). The RNF145 CDS, PCR amplified from an RNF145 IMAGE clone (Source Bioscience, Nottingham, UK), was cloned into pHRSIN-P_SFFV_-GFP-P_PGK_-Hygromycin^R^ (BamHI, NotI) (Demaison et al., 2002), replacing GFP with the transgene. To generate RNF145-V5, RNF145 CDS was Gibson cloned into pHRSIN-P_SFFV_-P_PGK_-Hygromycin^R^ containing a downstream in-frame V5 tag. RNF145-V5 RING domain mutations (C552A, H554A) were introduced by PCR amplification of RNF145-V5 fragments with primers encoding for C552A and H554A mutations and RNF145(C552A, H554A)-V5 was introduced into pHRSIN-P_SFFV_-P_PGK_-Hygromycin^R^ by Gibson assembly. FLAG-NLS-Cas9 was cloned from the lentiCRISPR v2 (Sanjana et al., 2014) (Addgene #49535, deposited by Feng Zhang) into pHRSIN.pSFFV MCS(+) pSV40 Blast (BamHI, NotI).

### Compounds

The following compounds were used in this study: Dulbecco’s Modified Eagle’s Medium high glucose (DMEM; Sigma-Aldrich, 6429-500ml), foetal calf serum (FCS; Seralab (catalogue no: EU-000, SLI batch: E8060012, Supplier batch: A5020012) and Life Technologies (catalogue no: 10270, lot: 42G4179K)), lipoprotein-deficient serum (LPDS; biosera, FB-1001L/100), mevastatin (Sigma-Aldrich, M2537-5MG), mevalonolactone (Sigma-Aldrich, M4467-1G), cholesterol (Sigma-Aldrich, C3045-5G), 25-hydroxycholesterol (Sigma-Aldrich, H1015-10MG), methyl-β-cyclodextrin (MBCD; Sigma-Aldrich, 332615-1G), bortezomib/PS-341 (BostonBiochem, I-200), (S)-MG132 (Cayman Chemicals, 10012628), NMS-873 (Selleckchem, s728501), digitonin (Merck, 300410-5GM), cycloheximide (Sigma-Aldrich, C-7698), IgG Sepharose™ 6 Fast Flow (GE Healthcare, 17-0969-01), ProLong™ Gold Antifade Mountant with DAPI (Thermo Fisher), bovine serum albumin (BSA; Sigma-Aldrich, A4503-10G), Protein A-Sepharose^R^ (P3391-1.5G), iodoacetamide (IAA; Sigma-Aldrich, I1149-5G), cOmplete protease inhibitor (EDTA-free; Roche, 27368400), phenylmethylsulfonyl fluoride (PMSF; Roche, 20039220), V5 peptide (Sigma-Aldrich, V7754-4MG), N-ethylmaleimide (NEM; Sigma-Aldrich, E3876-5G), puromycin (Cayman Chemicals, 13884), hygromycin B (Invitrogen, 10687010), Penicillin-Streptomycin (10,000 U/mL; Thermo Fisher, 15140122).

### Antibodies

Antibodies specific for the following targets were used for immunoblotting analysis: Insig-1 (rabbit; Abcam, ab70784), Hrd1 (rabbit; Abgent, AP2184a), TRC8 (rabbit; Santa Cruz, sc-68373), tubulin (mouse; Sigma, T9026), VCP (mouse; abcam, ab11433), β-actin (mouse; Sigma-Aldrich, A5316), calnexin (mouse; AF8, kind gift from M Brenner, Harvard Medical School), calreticulin (rabbit; Pierce, PA3-900), HMGCR (mouse; Santa Cruz, sc-27195), HMGCR (rabbit; Abcam, ab174830), gp78 (rabbit; ProteinTech, 16675-1-AP), Insig-1 (rabbit; Abcam, ab70784), RNF145 (rabbit; ProteinTech, 24524-I-AP), V5 (mouse; Abcam, ab27671), VU-1 ubiquitin (mouse; Life Sensors, VU101), UBE2G2 (mouse; Santa Cruz, sc-100613), GFP (rabbit; Life technologies, A11122), KDEL (mouse; Enzo, 10C3), HRP-conjugated anti-mouse and anti-rabbit (goat; Jackson ImmunoResearch), TrueBlot^®^ Anti-Rabbit-HRP (Rockland, 18-8816-31), TrueBlot^®^ Anti-Mouse-HRP ULTRA (Rockland, 18-8817-30). Alexa Fluor 488 (goat anti-rabbit; Thermo Fisher), Alexa Fluor 568 (goat antimouse; Thermo Fisher) were used as secondary antibodies for immunofluorescence microscopy. Anti-MHC-I (W6/32; mouse) and Alexa Fluor 647 (rabbit anti-mouse; Thermo Fisher) were used for cytofluorometric analysis.

### Cell Culture

HeLa, HEK-293T, Huh-7 and HepG2 cells were maintained in DMEM + 10% FCS + penicillin/streptomycin (1:100) (5% CO_2_, 37°C). Transfection of HeLa cells was performed using the TransIT-HeLa MONSTER kit (Mirus) according to the manufacturer’s instructions. Cells were seeded at low confluency in 12-well tissue culture plates and the next day transfection mix (1 μg DNA, 3 μl TransIT-HeLa reagent + 2 μl MONSTER reagent in OptiMEM (Gibco)) was added. Alternatively, reverse transfection was performed by seeding 3.5*10^5^ cells per well of a 12-well plate to the transfection mix on the day of transfection. For co-transfection of multiple plasmids, equal amounts of each plasmid were added up to 1 μg.

### CRISPR/Cas9-mediated gene knockout

CRISPR/Cas9-mediated genomic editing was performed according to Ran *et al*. (Ran et al., 2013). For generation of knockout cell lines, cells were transfected with pSpCas9(BB)-2A-Puro (PX459) V1.0 or V2.0 (Addgene #48139, and #62988 respectively; deposited by Dr. Feng Zhang) containing a sgRNA specific for the targeted gene of interest. Guide RNA sequences are listed in **Supplementary File 4**. Single cell clones were derived from cells transfected with a single sgRNA, whereas mixed knockout populations were generated by introducing 1 − 4 sgRNAs (**Supplementary File 5** for cell lines used in this study). Cells were cultured for an additional 24h before selection with puromycin (2 μg/ml) at low confluency for 72h. The resulting mixed knockout populations were used to generate singlecell clones by limiting dilution. Gene disruption was validated by immunoblotting, immunoprecipitation and/or targeted genomic sequencing.

### CRISPR/Cas9-mediated gene knock-in

An HMGCR-Clover knock-in donor template was created by Gibson assembly of ~ 1 kb flanking homology arms, PCR-amplified from HeLa genomic DNA, and the NsiI and PciI digested backbone from pMAX-GFP (Amaxa), into the loxP-Ub-Puro cassette from pDonor loxP Ub-Puro (kind gift from Prof Ron Kopito, Stanford University). Each arm was amplified using nested PCR. The 5’ arm was amplified using 5’-GATGCAGCACAGAATGTTGGTAG-3’ and 5’-CAATGCCCATGTTCCAGTTCAG-3’, followed by 5’-CAATGCCCATGTTCCAGTTCAG-3’ and 5’-CAGCTGCACCATGCCATCTATAG-3’. The 3’ arm was amplified using the following primer pairs: 5’-CCAAGGAGCTTGCACCAAGAAG-3’ and 5’-CTAAGGTCCCAGTCTTGCTTG-3’. The product served as template for a subsequent PCR step using the primers 5’-CCAAGGAGCTTGCACCAAGAAG-3’ and 5’-GTCACCCTCATCTAAGCAAC-3’. Overhangs required for Gibson assembly were introduced by PCR. HeLa cells were co-transfected with Cas9, sgRNA targeting immediately downstream of the HMGCR stop codon and donor template. Three different donor templates were simultaneously transfected, each differing in the drug resistance marker (puromycin, hygromycin and blasticidin). The transfected cells were treated with the three antibiotics five days post-transfection until only drug-resistant cells remained. The resulting population was transfected with Cre-recombinase in pHRSIN MCS(+) IRES mCherry pGK Hygro. mCherry positive cells were single-cell cloned by FACS.

### Lentivirus production and transductions

HEK-293T cells were transfected with a lentiviral expression vector, the packaging vectors pCMVΔR8.91 and pMD.G at a ratio of 1:0.7:0.3 using TransIT-293 (Mirus) as recommended by the manufacturer. For production of CRISPR library virus, HEK-293T cells were transfected as above in 15 cm tissue culture plates. 48 h post transfection, virus-containing media was collected, filtered (0.45 μm pore size) and directly added to target cells or frozen (-80°C) for long-term storage. Cells were transduced in 6-well tissue culture plates at an M.O.I. < 1 and selected with puromycin (2 μg/ml) or hygromycin B (200 μg/ml). To generate HeLa HMGCR-Clover stably expressing Cas9, HeLa HMGCR-Clover cells were transduced with pHRSIN-P_SFFV_-Cas9-P_PGK_-Hygromycin^R^ and stable integrants selected with hygromycin B. Cas9 activity was confirmed by transduction with pKLV encoding a β-2-microglobulin (B2M)-targeting sgRNA followed by puromycin selection. MHC-I surface expression was assessed by flow cytometry in puromycin-resistant cells five days post transduction. Typically, ~ 90% reduction of cell surface MHC-I expression was observed.

### Fluorescent PCR

To identify CRISPR-induced frame-shift mutations, genomic DNA was extracted from wild type HeLa cells and RNF145 CRISPR clones using the Quick-gDNA MicroPrep kit (Zymo Research) followed by nested PCR of the genomic region 5’ and 3’ of the predicted sgRNA binding site. One in each primer pair for the second PCR was 5’ modified with 6-FAM™ (fluorescein, Sigma-Aldrich). Primer sequences were as follows: For sgRNA #8 PCR1_Forward: CAGAATGCTCACTAGAAGATTAG, PCR1_Reverse: GTAGTATACGTTCTCACATAG, PCR2_Forward: GTGATGTAGACACTCACCTAC and PCR2_Reverse GTGACAACCTATTAGATTCGTG. PCR products were detected using an ABI 3730xl DNA Analyser.

### Flow cytometry and Fluorescence-activated cell sorting (FACS)

Cells were collected by trypsinisation and analysed using a FACS Calibur (BD) or an LSR Fortessa (BD). Flow cytometry data was analysed using the FlowJo software package. Cells resuspended in sorting buffer (PBS + 10 mM HEPES + 2% FCS) were filtered through a 50 μm filter, and sorted on an Influx machine (BD), or, for the ubiquitome CRISPR/Cas9 screen, on a FACS Melody (BD). Sorted cells were collected in DMEM + 50% FCS and subsequently cultured in DMEM + 10% FCS + penicillin/streptomycin. For MHC-I flow cytometric analysis, cells resuspended in cold PBS were incubated with W6/32 (20 min, 4°C), washed twice and then incubated with Alexa-647-labelled anti-mouse antibody (15 min, 4°C). Cells were washed twice and resuspended in PBS.

### CRISPR/Cas9 knockout screens

For genome-wide and ubiquitome CRISPR/Cas9 knockout screens, 10^8^ and 1.2*10^7^ HeLa HMGCR-Clover (Cas9) or ΔRNF145 #6 (Cas9), respectively, were transduced at M.O.I. ~ 0.3 by spinfection (750xg, 60 min, 37°C). Transduction efficiency was determined *via* flow-cytometry-based measurement of BFP expression 48-72h post infection. Transduced cells were enriched by puromycin (1 μg/ml). On day 9 (genome-wide screen) or day 7 (ubiquitome-library screen) post transduction, cells were rinsed extensively with PBS and cultured overnight in starvation medium (DMEM + 10% LPDS + 10 μM mevastatin + penicillin/streptomycin) before sterol addition (2 μg/ml 25-hydroxycholesterol and 20 μg/ml cholesterol for 5h). An initial FACS selection (‘sort 1’) on cells expressing high levels (~0.3-0.6% of overall population) of HMGCR-Clover (HMGCR-Clover^high^) was performed. 2*10^5^ (genome-wide screen) and ~ 10^5^ (ubiquitome screen) sorted cells were pelleted and DNA was extracted using the Quick-gDNA MicroPrep kit (Zymo Research). To gauge sgRNA enrichment, DNA was extracted from 3*10^6^ (genome-wide library screen) or 6*10^6^ (ubiquitome library screen) cells pre-sort using the Gentra Puregene Core kit A (Qiagen). Cells in the genome-wide screen were subjected to a second round of sterol deprivation and sort (see above) after expansion of initially 2.5*10^5^ sorted cells for 8 days. Sorted cells were cultured until 5*10^6^ cells could be harvested for genomic DNA extraction using the Gentra Puregene Core kit A (Qiagen). Individual integrated sgRNA sequences were amplified by two sequential rounds of PCR, the latter introducing adaptors for Illumina sequencing (Supplementary File 3). Sequencing was carried out using the Illumina HighSeq (genome-wide screen) and MiniSeq (ubiquitome screen) platforms. Illumina HiSeq data was analysed as described previously (Timms et al., 2016). Guide RNA counts were analysed with the RSA algorithm under default settings (König et al., 2007). Of note, a gene’s calculated high significance value and therefore high enrichment in the selected population does not necessarily reflect its importance relative to genes with lower significance values/enrichment, since gene disruption can be incomplete or lethal phenotypes might evade enrichment.

### Quantitative PCR

Whole-cell RNA was isolated with the RNeasy Plus Mini Kit (Qiagen, Venlo, Netherlands) and reverse transcribed using Oligo(dT)15 primer (Promega, C110A) and SuperScript™ III reverse transcriptase (Invitrogen). Transcript levels were determined in triplicate using SYBR^®^ Green PCR Master Mix (Applied Biosystems) in a real time PCR thermocycler (7500 Real Time PCR System, Applied Biosystems). Primers used for target amplification can be found in **Supplementary file 2**. RNA quantification was performed using the ΔΔCT method. GAPDH transcript levels were used for normalization.

### Sterol depletion assays

HeLa cells at ~ 50% confluency were washed five times with PBS and cultured for 16-20h in starvation medium (DMEM + 10% LPDS + 10 μM mevastatin + penicillin/streptomycin) before addition of 25-hydroxycholesterol (2 μg/ml) and cholesterol (20 μg/ml) to analyse sterol-accelerated protein degradation. For experiments in **Figures 5C, D and 6**, sterol-depletion was performed in starvation medium + 50 μM mevalonate.

### Chol:MBCD complex preparation

Complexation of cholesterol (2.5 mM) with MBCD (25 mM) was performed according to Christian *et al*. (Christian et al., 1997). An emulsion of cholesterol powder (final: 2.5 mM) and an MBCD solution (25 mM) was produced by vortexing and tip sonication (1 min in 10 s intervals), and continuously mixed for 16h at 37°C. The solution was sterile filtered (0.45 μm PVDF pore size) and stored at -20°C.

### Preparation of sterols and mevalonate

Sterols were prepared by resuspension in ethanol or complexation with MBCD (see above). Mevalonate was prepared by adding 385 μl 2.04M KOH to 100 mg mevalonolactone (Sigma). The solution was heated (1h, 37°C) and adjusted to a 50 mM stock solution.

### SDS-PAGE and immunoblotting

Cells were collected mechanically in cold PBS or by trypsinisation, centrifuged (1000xg, 4 min, 4°C), and cell pellets resuspended in lysis buffer (1% (w/v) digitonin, 1x cOmplete protease inhibitor, 0.5 mM PMSF, 10 mM IAA, 2 mM NEM, 10 mM TRIS, 150 mM NaCl, ph 7.4). After 40 min incubation on ice, lysates were centrifuged (17.000xg, 15 min, 4°C), the post-nuclear fraction isolated and protein concentration determined by Bradford assay. Samples were adjusted with lysis buffer and 6 x Laemmli buffer + 100 mM dithiothreitol (DTT) and heated at 50°C (15 min). Samples were separated by SDS-PAGE and transferred to PVDF membranes (Merck) for immunodetection. Membranes were blocked in 5% milk + PBST (PBS + 0.2% (v/v) Tween-20) (1h) and incubated with primary antibody in PBST + 2% (w/v) BSA at 4°C overnight. For detection from whole-cell lysate, membranes were incubated in peroxidase (HRP)-conjugated secondary antibodies. For detection of immunoprecipitated proteins, TrueBlot^®^ HRP-conjugated secondary antibodies (Rockland) were used. Immunoprecipitated RNF145 was detected using Protein A-conjugated HRP.

### Immunoprecipitation

Cells were seeded to 15 cm tissue culture plates (4*10^6^ cells per plate). The following day, cells were washed five times with PBS and cultured in starvation medium (DMEM + 10% LPDS + 10 μM mevastatin + 10 μM mevalonate + penicillin/streptomycin) for 20h. To prevent HMGCR membrane extraction and degradation, starved cells were treated with NMS-873 (50 μM) 0.5h prior to sterol addition (2 μg/ml 25-hydroxycholesterol and 20 μg/ml cholesterol for 1h) and collection in cold PBS. Cells were lysed in IP buffer 1 (1% (w/v) digitonin, 10 μM ZnCl_2_, 1x cOmplete protease inhibitor, 0.5 mM PMSF, 10 mM IAA, 2 mM NEM, 10 mM TRIS, 150 mM NaCl, ph 7.4), post-nuclear fractions isolated by centrifugation (17.000×g, 4°C, 15 min) adjusted to 0.5% (w/v) digitonin and pre-cleared with IgG Sepharose™ 6 Fast Flow (1h). Endogenous RNF145 and V5-tagged RNF145 were immunoprecipitated at 4°C overnight from 3 - 6 mg whole-cell lysate using Protein A-Sepharose^R^ and anti-RNF145 or V5 antibody, respectively. Beads were collected by centrifugation (1500xg, 4 min, 4°C), washed for 5 min with IP buffer 2 (0.5% (w/v) digitonin, 10 μM ZnCl_2_, 10 mM Tris, 150 mM NaCl, ph 7.4) and 4 x 5 min with IP buffer 3 (0.1% (w/v) digitonin, 10 μM ZnCl_2_, 10 mM TRIS, 150 mM NaCl, ph 7.4). Proteins whose interaction with RNF145 was labile in the presence of 1% (v/v) Triton X-100 were recovered by eluting twice with 20 μl TX100 elution buffer (1% (v/v) Triton X-100 + 2x cOmplete protease inhibitor in 10 mM TRIS, 150 mM NaCl pH 7.4) at 37°C under constant agitation. Immunoprecipitated RNF145 was subsequently eluted in 30 μl 2x Laemmli buffer + 3% (w/v) DTT. RNF145-V5 and associated complexes were recovered by two sequential elutions with V5 elution buffer (1 mg/ml V5 peptide + 2x cOmplete protease inhibitor in 10 mM TRIS, 150 mM NaCl pH 7.4) for 30 min under continuous agitation. Eluted samples were adjusted Laemmli buffer and denatured at 50°C for15 min.

### Ubiquitination assays

Cells were sterol-depleted (20h), treated with 20 μM MG132 and left for 30 min before addition of sterols (2 μg/ml 25-hydroxycholesterol and 20 μg/ml cholesterol for 1h) or EtOH (vehicle control). Immunoprecipitation of ubiquitinated HMGCR was performed as described above from 1 mg whole-cell lysate and using rabbit α-HMGCR (Abcam, ab174830). Proteins were eluted in 30 μl 2x Laemmli buffer + 100 mM DTT at 50°C for 15 min. For immunoblotting of ubiquitin with mouse VU-1 α-ubiquitin (Life Sensors, VU101), the PVDF membrane was incubated with 0.5% (v/v) glutaraldehyde/PBS pH 7.0 (20 min) and washed 3x with PBS prior to blocking in 5% (w/v) milk + PBS + 0.1% (v/v) Tween-20.

### Indirect immunofluorescence confocal microscopy

Cells were grown on coverslips, fixed in 4% PFA (15 min), permeabilised in 0.2% (v/v) Triton X-100 (5 min) and blocked with 3% (w/v) BSA/PBS (30 min). Cells were stained with primary antibody diluted in 3% (w/v) BSA/PBS (1h), washed with 0.1% (w/v) BSA/PBS, followed by staining with secondary antibody in 3% BSA/PBS (1h), an additional washing step (0.1% (w/v) BSA/PBS) and embedded using ProLong™ Gold Antifade Mountant with DAPI (Thermo Fisher). Images were acquired using an LSM880 confocal microscope (Zeiss) at 64x magnification.

### Statistical analysis

Statistical significance was calculated using the unpaired Student’s t-test.

### Data deposition

Sequencing data from CRISPR/Cas9 knockout screens presented in this study have been deposited at SRA (genome-wide screen: SUB4198636; ubiquitome screen: SUB4183663).

## SUPPLEMENTARY FILES

**Supplementary File 1**. sgRNA sequences and genes targeted by the CRISPR/Cas9 ubiquitome library.

**Supplementary File 2**. Primer sequences used for qPCR.

**Supplementary File 3**. Primers used in CRISPR/Cas9 screens.

**Supplementary File 4**. sgRNA sequences for generation of knockout cell lines.

**Supplementary File 5**. Genetically modified cell lines used in this study.

